# Constructing and Optimizing 3D Atlases From 2D Data With Application to the Developing Mouse Brain

**DOI:** 10.1101/2020.04.01.017665

**Authors:** David M Young, Siavash Fazel Darbandi, Grace Schwartz, Zachary Bonzell, Deniz Yuruk, Mai Nojima, Laurent Gole, John Rubenstein, Weimiao Yu, Stephan J Sanders

**Affiliations:** Department of Psychiatry, UCSF Weill Institute for Neurosciences, University of California, San Francisco, San Francisco, California 94158, USA; Institute of Molecular and Cellular Biology, Agency for Science, Technology and Research, Singapore 138673, Singapore

## Abstract

3D imaging data necessitate 3D reference atlases for accurate quantitative interpretation. Existing computational methods to generate 3D atlases from 2D-derived atlases result in extensive artifacts, while manual curation approaches are labor-intensive. We present a computational approach for 3D atlas construction that substantially reduces artifacts by identifying anatomical boundaries in the underlying imaging data and using these to guide 3D transformation. Anatomical boundaries also allow extension of atlases to complete edge regions. Applying these methods to the eight developmental stages in the Allen Developing Mouse Brain Atlas (ADMBA) led to more comprehensive and accurate atlases. We generated imaging data from fifteen whole mouse brains to validate atlas performance and observed qualitative and quantitative improvement (37% greater alignment between atlas and anatomical boundaries). We provide the methods as the MagellanMapper software and the eight 3D reconstructed ADMBA atlases. These resources facilitate whole-organ quantitative analysis between samples and across development.

## Introduction

Anatomical atlases have played a crucial role in research into organogenesis, anatomy, physiology, and pathology [1]–[3] and have been used for a wide range of applications, including investigating developmental biology [4], defining stereotactic coordinates [5], identifying patterns of neural activity associated with behavior [6], [7], and integrating morphological, transcriptomic, and neural activity data across species [1]. Ongoing initiatives continue to develop these resources, including the Human Cell Atlas [8], the BRAIN Initiative [9], [10], the Human Brain Project [11], and centralized repositories of data, including the Allen Brain Atlas [12]–[14].

Macroscale imaging, such as magnetic resonance imaging (MRI), routinely generates data from intact whole organs [15]–[21] in contrast to microscale imaging, such as light microscopy, which has typically relied on physically cutting thin sections leading to misalignment artifacts. Technological advances in microscopy have facilitated a dramatic increase in imaging of high-resolution, intact whole organs using serial twophoton tomography (STPT) [22] or tissue clearing techniques (e.g. CLARITY [23], 3DISCO [24], and CUBIC [25]) and lightsheet microscopy [26]–[31]. These 3D microscopy methods have been applied to map cytoarchitecture and cellular connectivity in complex tissues, including the brain [30], [32], [33], heart [34], [35], and lung [36], [37].

Accurate analysis of imaging data from whole organs requires atlases of the corresponding organ, species, developmental age, resolution, and dimensions [38]–[41]. The progression to 3D microscopy data necessitates 3D anatomical atlases to reference data to existing anatomically-defined labels [42], [43]. Though numerous atlases based on 2D physical sections exist [5], [37], [44]–[46], most suffer from label misalignments or insufficient resolution in the third dimension, limiting their utility for 3D imaging [47]. The most recent version of the Allen Common Coordinate Framework atlas (CCFv3) of the adult (P56) C57BL/6J mouse brain represents one of the most complete 3D atlases at microscopic resolution to date [47]. A team of neuroanatomists and illustrators redrew structures natively in 3D, sometimes using the original annotations as seeds to guide the redrawing process, producing 658 structural annotations [47]. Even with this monumental effort, 23% of labels from the original Allen Reference Atlas were not included in the 3D version, largely due to the time and labor constraints of further manual curation [47]. To date, this titanic effort to generate a 3D atlas at microscopic resolution has not been repeated for most combinations of organ, species, and developmental stage, despite the numerous existing 2D-based atlases.

Automated methods to convert existed 2D-derived atlases into fully 3D atlases could leverage existing, established atlases without the time-consuming effort required for manual 3D curation, or provide the substrate for more rapid manual fine-tuning. This conversion needs to address multiple limitations in atlases based on 2D physical sections, as demonstrated by the challenges of using the existing 3D reconstruction of eight mouse brains, each at a different developmental time point, in the Allen Developing Mouse Brain Atlas (ADMBA) [46] (Fig. 1A). First, all atlases in the ADMBA are missing labels across large regions, typically the lateral planes of one hemisphere and the entire opposite hemisphere. Second, warping and asymmetrical placement of samples in the histological images prevents simply mirroring labels from the annotated to the non-annotated hemisphere. Third, as noted in the original Allen Reference Atlas [14], [40], [47], the assembled 3D volume is smooth in the sagittal planes, in which the atlas annotations were drawn, but suffers from substantial edge artifacts in the coronal and axial planes (Fig. 1A).

**Figure 1:**
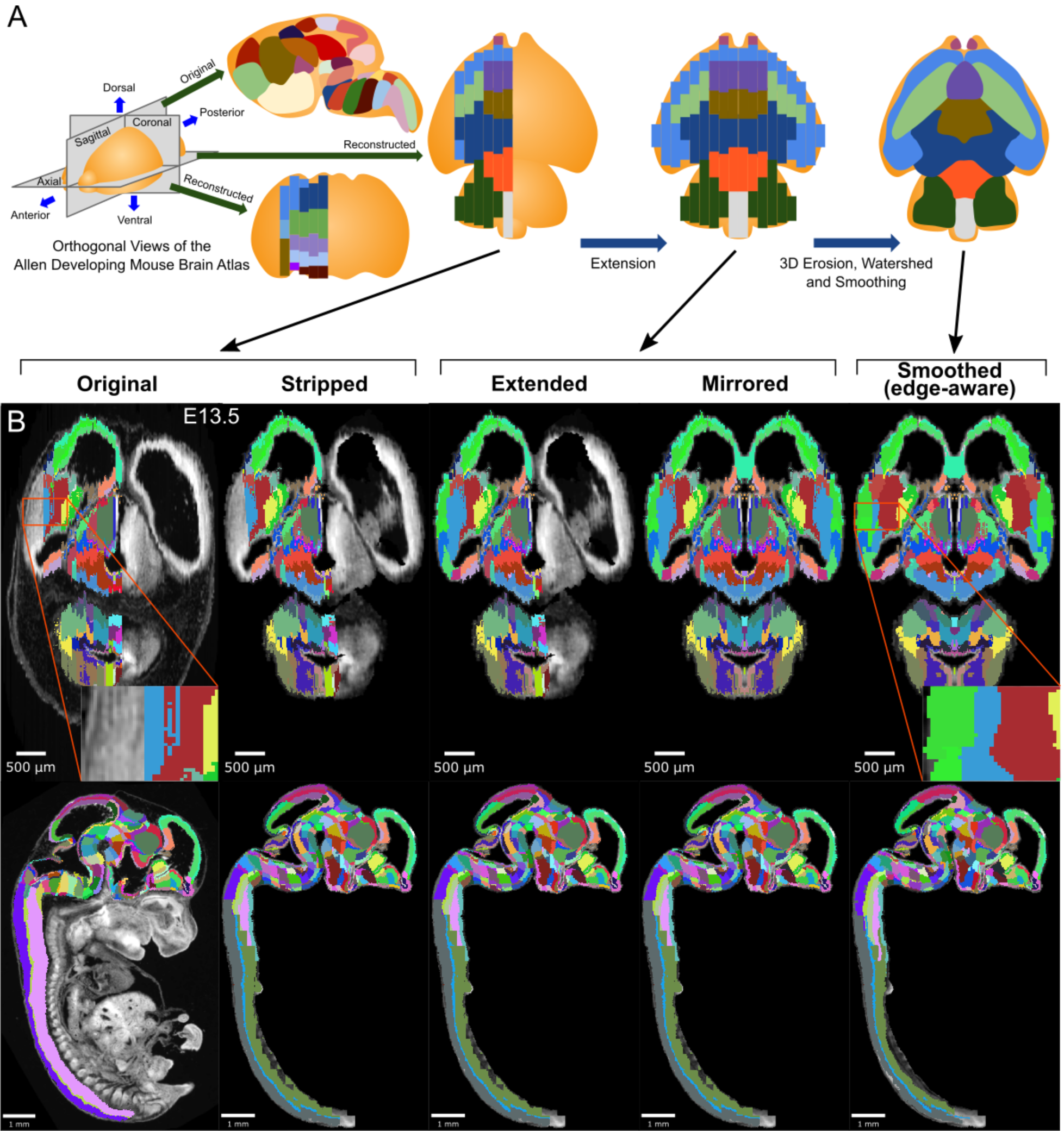
3D Atlas Refinement Pipeline Overview. (A) 2D-derived atlases, such as those in the Allen Developing Mouse Brain Atlas, are smooth and consistent in the sagittal plane in which they were annotated. However, in the 3D reconstructions of these 2D sagittal planes, the coronal and axial planes reveal missing sections and jagged edges. To improve their performance for annotating 3D data, the lateral edges are extended to complete the labeled hemisphere. A 3D rotation is applied to bring the brain parallel to the image borders, then both the completed hemisphere labels and underlying microscopy sections are mirrored across the sagittal midline to complete coverage in the opposite hemisphere. To improve anatomical registration, the labels are each eroded and re-grown through a 3D watershed, guided by the anatomical edge map. To smooth the final product, labels are iteratively filtered with a morphological opening operation, or a closing operation for small, disconnected labels that would otherwise be lost. (B) The pipeline illustrated in axial (top) and sagittal (bottom) views on the ADMBA E13.5 atlas, which requires the full pipeline shown in “A”, including an additional step to strip out non-CNS tissue from the original whole embryo imaging. The nomenclature for pipeline steps shown here is used consistently throughout the manuscript. Insets of the “Original” and “Smoothed (edge-aware)” lateral regions highlight the label extension and smoothing. A spreadsheet mapping colors to label names and IDs in the Allen ontology for each atlas in this manuscript can be found at Data S1.

Several automated approaches have partially addressed some of these issues. The Waxholm Space rat brain atlas used two standard reference atlases as templates to yield an atlas with relatively smooth 3D structures through manual and semi-automated segmentation, though some label edge artifacts remain [48]. Ero, et al. [49] performed non-rigid alignments between histological slice planes of the Common Coordinate Framework (CCFv2, 2011), applying the same deformations to the label images. This registration improved alignment, however the labels often appear to extend beyond the underlying histological planes. Rather than registering planes, Niedworok, et al. [40] applied a smoothing filter on the annotations themselves to reduce label artifacts, though many label edge irregularities persisted.

Here, we present a method to automatically generate fully 3D atlases from existing 2D-derived partial atlases and apply it to the full ADMBA. We performed three steps (Fig. 1A): 1) Extension and mirroring of 2D data to approximate missing labels, 2) Smoothing with 3D watershed and morphological methods to align jagged edges between 2D sections, and 3) Edge detection from microscopy images to guide the 3D watershed to align boundaries with anatomical divisions, which we call “edge-aware” refinement (example atlas shown in Fig. 1B). We demonstrate qualitative and quantitative improvements over the existing 3D ADMBA. To assess the resulting 3D atlas, we generated 3D imaging data from 15 C57BL/6J wild-type mouse brains at postnatal day 0 (P0) using tissue clearing and lightsheet microscopy. The 3D labels in our refined atlas match the brain structures better than the initial atlas, as demonstrated by closer alignment between labels and anatomical edges and decreased variability within labels at the cellular level. We provide the method as open source software called MagellanMapper to apply the algorithms to equivalent atlases and provide the resulting ADMBA 3D atlases as a resource for immediate application to automated whole-brain 3D microscopy analysis.

## Results

### Allen Developmental Mouse Brain Atlas

The ADMBA contains data for eight mouse brains each at a different development stage: embryonic day 11.5, E13.5, E15.5, E18.5, postnatal day (P) 4, P14, P28, and P56. The data are generated from 158 to 456 serial 2D sagittal sections of 20-25 *µm* stained with Nissl or Feulgen-HP yellow and imaged with a light microscope at a lateral resolution of 0.99 to 1.049 *µm*. For the three earliest stages of development, E11.5, E13.5, E15.5, the entire embryo was imaged, rather than just the brain. Each of the eight atlases are annotated with expertly-curated labels of brain structures [46], [50] in a hierarchy starting with the largest structures (e.g. neural plate) extending through 13 sublevels to the smallest annotated structures (e.g. parvicellular part of Lat). Viewed sagittally, these labels cover the majority of tissue in each brain with smooth, anatomically-aligned edges (Fig. 2A), however only sections on the left side of each brain are annotated and, for six atlases, labels are not present for the most lateral 14-24% sagittal planes of the brain (4-5% by volume; Suppl. Fig. S2; Fig. 2B). Labels extend slightly beyond the midline for several atlases, helping to annotate brains with a midline skew, yielding a median label to atlas volume ratio of 51%.

**Figure 2:**
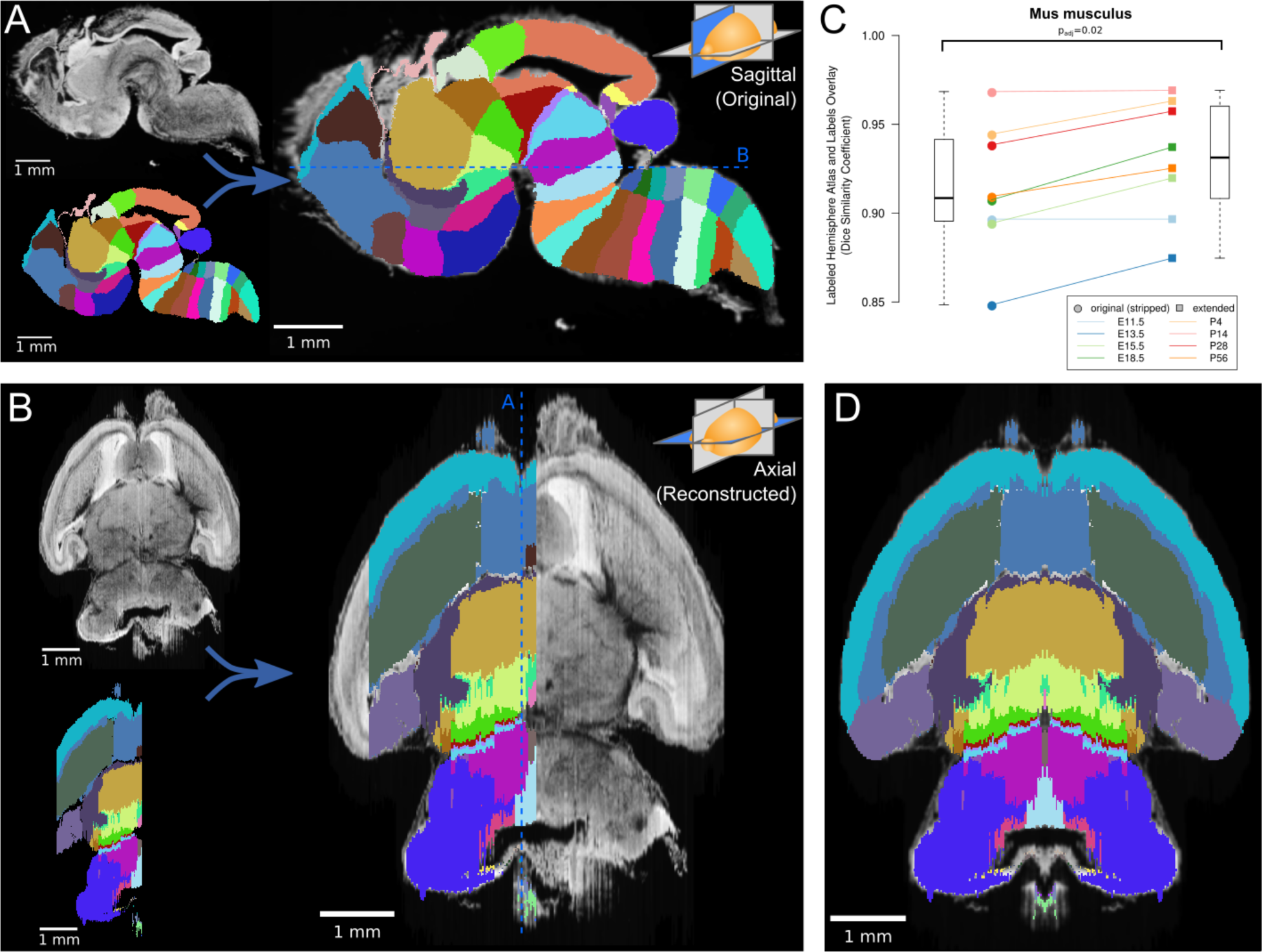
Atlas extension. (A) The original E18.5 atlas labels (bottom left) viewed sagittally demonstrate smooth borders and close correspondence with the underlying microscopy images (upper left; overlaid on right). The dashed blue line shows the section viewed in “B”. (B) When viewed axially, the most lateral sections lack labels, and label borders are jagged. Slight rotation of the underlying microscopy images leads to asymmetry between the two hemispheres. (C) The Dice Similarity Coefficient (DSC), a measure of the completeness of labeling compared to the thresholded atlases, for the labeled hemispheres increased for all brains in the ADMBA after lateral edge extension (original median = 0.91, extended median = 0.93; p = 0.02, Wilcoxon signed-rank test (WSRT); “Mus musculus,” level -1, ID 15564 in the Allen ontology). (D) To fill in the lateral edges using existing labels, a representative lateral label plane was iteratively resized to fit the underlying microscopy images. The plane for each subsequent microscopy plane was thresholded, the bounding box extracted, and the labels resized to fit this bounding box, followed by conforming labels to the underlying gross anatomical boundaries (Suppl. Fig. S1). A stretch of compressed planes was expanded (Suppl. Fig. S7), and the completed hemisphere of labels mirrored to the other hemisphere after rotation for symmetry to complete the labeling.

### Atlas similarity

To focus our analysis on the brain tissue, we excluded non-central nervous system (CNS) tissue for the three embryos for which the brain was not dissected prior to imaging. Across all eight atlases the similarity between the microscopy images and the original label annotations was estimated by taking the threshold of each sagittal section on the left hemisphere of the brain as an approximation of ground truth and comparing the label coverage using the Dice Similarity Coefficient (DSC) [51], calculated by Insight Segmentation and Registration Toolkit (ITK) [52]. Higher DSCs reflect greater similarity between the images and labels, with a maximum possible value of 1. The observed DSCs for the original labels ranged from 0.85 to 0.97 (median 0.91; Fig. 2).

### Atlas extension

To generate labels across the entire brain, we extended the existing labels to the lateral edges by following histological boundaries before mirroring the labels on the opposite hemisphere (Fig. 2). The most lateral labeled sagittal section provided the initial seed from which to grow labels laterally. First, we resized this plane to match the extent of corresponding microscopy signal in the next, unlabeled lateral sagittal section (Suppl. Fig. S1A, B). Next, we refined label boundaries by eroding each label and regrowing it along histological boundaries, which we call “edge-aware” refinement. To model these boundaries, we generated gross anatomical maps in 3D using edge-detection methods on the volumetric histology images. Taking the Laplacian of Gaussian [53] of each histological volumetric image, using a relatively large Gaussian sigma of 5, highlighted broad regions of similar intensities. A zero-crossing detector converted this regional map into a binary anatomical edge map (Suppl. Fig. S1B). After eroding each label, we next used this anatomical map to guide the regrowth of labels by a compact watershed algorithm [54] step, which adds a size constraint to the classic watershed algorithm. Distances of each pixel to its nearest edge map pixel formed anatomically based watershed catchment areas, while each eroded label served as a large seed from which filling of the nearest catchment areas began and grew until meeting neighboring labels. Thus, we extended a labeled sagittal plane to an unlabeled one, refined by the histology.

This process was repeated iteratively across all remaining lateral sections (Suppl. Fig. S1B, C). By using a 3D anatomical map for the watershed and seeding it with the prior plane’s labels, we ensured continuity between planes. To model the tapering of labels laterally, we preferentially eroded central labels by weighting each label’s filter size based on the label’s median distance from the tissue section’s outer boundaries. Central labels thus eroded away faster, while the next most central labels grew preferentially to fill these vacated spaces. In the E18.5 atlas, for example, this approach allowed the basal ganglia to taper off and give way to the dorsal and medial pallium (Fig. 2B). Although edges between planes remained somewhat jagged, this extension step creates a substrate for further refinement.

After completing label coverage for the left hemisphere, both the labels and underlying microscopy images were reflected across the sagittal midline to cover the remaining hemisphere. Care was taken to ensure that the sagittal midline was identified correctly by inspecting and rotating the 3D image and midline plane from multiple angles (Fig. 2C). Recalculating the DSC between the microscopy images and labels for the left hemisphere showed greater similarity across all eight atlases with a median DSC improvement of 0.02 (p = 0.02, WSRT) and a resulting DSC range of 0.87 to 0.97 (median 0.93, Fig. 2D). Equivalent analysis of DSC for the whole brain would show substantial improvement due to the absence of labels on the right side in the original.

### Label smoothing

The ADMBA atlases have been provided as 3D volumetric images, combined computationally from the original 2D sagittal reference plates; 2D sections can be generated from these 3D images in orthogonal dimensions to the original sagittal view. Visual inspection of labels in the axial and coronal planes reveals high-frequency artifacts along most edge borders, likely from the difficulty of drawing contiguous borders in dimensions orthogonal to the drawing plane (Figs. 1-3). To quantify the degree of label smoothness, we used the unit-less compactness metric [55]. The compactness measure applied in 3D incorporates both surface area and volume, allowing for quantification of smoothness to measure shape irregularity. Of note, compactness is independent of scale or orientation, facilitating comparison across all labels, despite size differences, and its sensitivity to noise allows finer detection of label irregularity [56]. Measuring the compactness of each label and taking the weighted mean based on label volume for each atlas gave a median compactness of 13,281 (mean: 33,975, standard deviation (SD): 44,743). For context, across all eight atlases the 3D whole-brain microscopy images were more compact (median compactness: 1,895, mean: 2,615, SD: 2,305; p = 0.02, WSRT, Bonferroni corrected; Suppl. Fig. S3A), consistent with the observed irregularity in the label images compared to anatomical images (Figs. 2B and 3).

To reduce this irregularity, we applied a smoothing filter iteratively to the 3D image of each label. Prior approaches to this problem used a Gaussian filter (kernel SD of 0.5, applied in two passes) [40]. While this visually improved the smoothness of label edges, we observed sharp angles along the edges, presumably from the limitation of rounding blurred pixel values to the integers required for label identities rather than allowing the subtler floating-point gradients from the Gaussian kernel (Suppl. Fig. S4A). In addition, the Gaussian filter expanded the volume of each label, leading to label loss as larger labels enveloped smaller ones (Suppl. Fig. S4B).

Optimal smoothing would maximize the smoothness of each label (i.e. compactness [56]) whilst minimizing changes in shape and location. It would also not lead to label loss. To calculate the improvement in compactness, we defined “compaction” as the difference in compactness between the original and smoothed label over the compactness of the original label with a range from 0 or 1, with 1 being optimal. For changes in shape and location, we defined “displacement” as the fraction of the smoothed label that was outside of the original label with a range from 0 or 1, with 0 being optimal. We defined a “smoothing quality” metric to reflect the balance of compaction and displacement, calculated as the difference between these two measures with a range from -1 or 1, with 1 being optimal. To estimate atlas-wide smoothing quality, we took the weighted sum by volume of smoothing quality for all labels in the atlas. Assessing the quality of smoothing using Gaussian blur with increasing Gaussian sigmas, we observed label loss in all atlases in the ADMBA, even with a small sigma where labels remained visibly jagged. At this sigma value, the median atlas-wide smoothing quality across all eight atlases was 0.54 (mean 0.53), rising to a peak of 0.57 (mean 0.56) at sigma 0.5 and 0.49 (mean 0.53) at a sigma of 1.0, but with substantial loss of labels (median 14% lost at sigma 0.25, rising to 42% lost at sigma 1; mean 11% and 47%, respectively Suppl. Fig. S4B).

To refine smoothing while minimizing label loss, we changed the filter from a Gaussian to a morphological opening filter [57]. This filter first erodes each label to remove artifacts, followed by dilation to restore its original volume. To avoid label loss caused by excessive erosion of small labels, we halved the size of its structuring element for labels with ≤5,000 pixels. A few labels were split into numerous tiny fragments that would disappear with an opening filter. For these small labels, the opening filter was replaced by a closing filter, reversing the process by dilating before eroding to reconnect these components. With this adaptive opening filter approach, labels became more compact with smoother edges, while retaining their overall shape as seen in both 2D and 3D visualizations (Fig. 3A).

**Figure 3:**
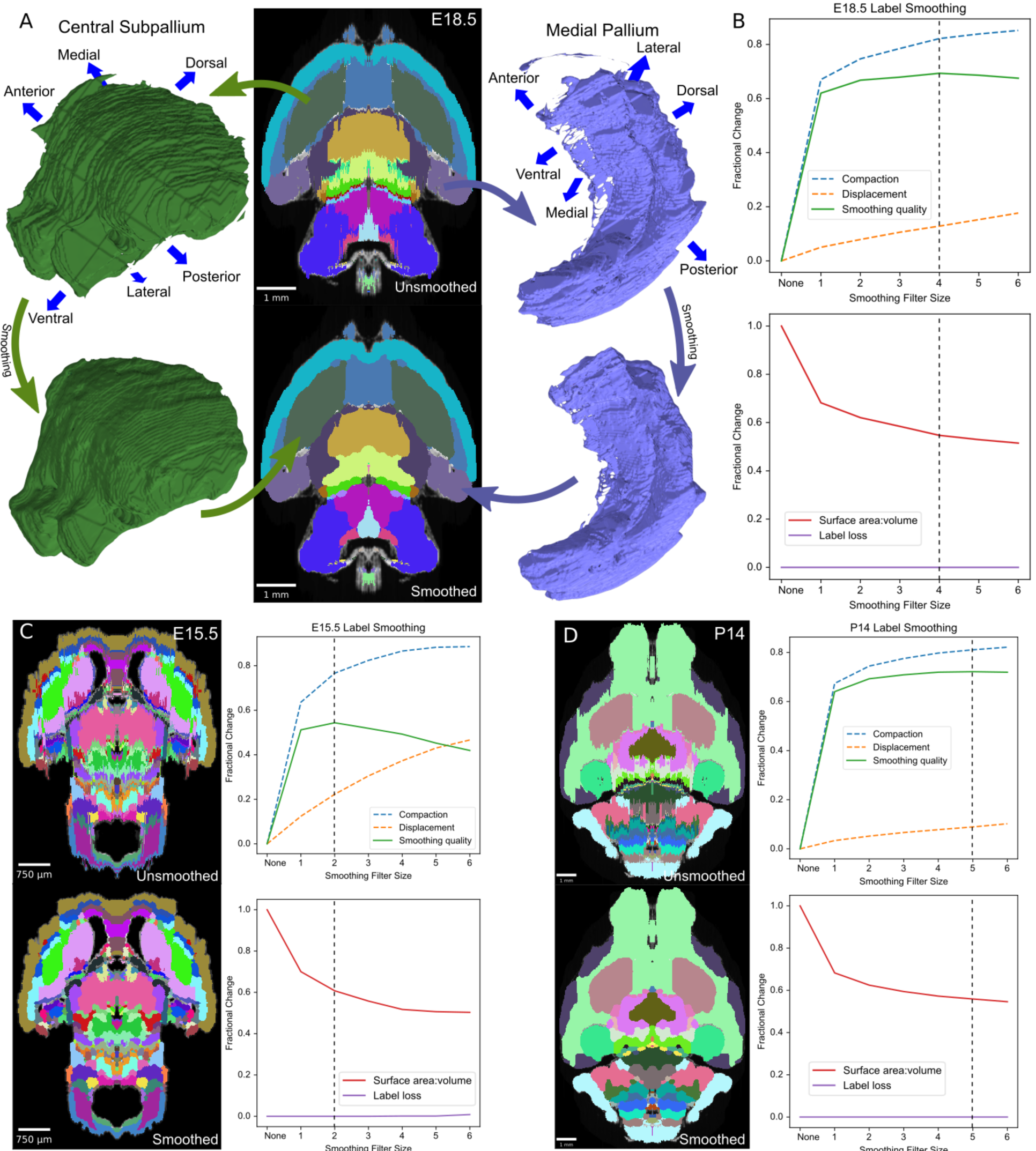
Atlas extension. (A) Irregular edges at the label edges in the original (mirrored) E18.5 atlas (top) were smoothed by applying an adaptive morphological opening filter iteratively to each label, starting from the largest to the smallest labels (bottom). 3D renderings are depicted for two representative labels before (top) and after (bottom) smoothing. (B) To identify the optimal filter structuring element size, we devised an “atlas-wide smoothing quality” metric to incorporate the balance between smoothing (compaction) and changes in size and shape (displacement). While compaction continued to improve with increasing structuring element size, displacement eventually caught up, giving an optimal atlas-wide smoothing quality with a structuring element size of 4 for the E18.5 atlas (top). We also assessed the number of labels that were lost (none in this example) and the surface area to volume ratio, for which lower values reflect smoother shapes (bottom). Vertical dashed lines indicate the optimal filter structuring element size, based on “atlas-wide smoothing quality”. (C) Selection of the optimal filter structuring element size using the same metrics as in “B” are shown for the E15.5 and (D) P14 atlases.

Quantifying the improvement with the adaptive opening filter approach, using only filter sizes that completely eliminated label loss, we obtained a median atlas-wide smoothing quality of 0.61 (mean 0.62) across all eight atlases, and improvement over the Gaussian filter approach (sigma 0.25; p = 0.008; Suppl. Fig. S4C). The optimal filter size varied between atlases, ranging from 2 to 5 (E15.5 and P14 shown in Fig. 3C, D; all ADMBA shown in Suppl. Fig. S5). The median overall compactness improved significantly (13,281 (SD = 44,743) for unsmoothed labels vs. 2,540 (SD = 2,624) for smoothed labels, p = 0.02, WSRT, Bonferroni corrected; mean 33,975 vs. 3,190) to a level that did not differ from that observed for the microscopy images of whole brains (p = 1.00, WSRT, Bonferroni corrected; Suppl. Fig. S3B, C).

Because morphological filters such as erosion classically operate globally, a drawback to these filters is the potential loss of thin structures, such as loss of the thin portion of the alar plate of the evaginated telencephalic vesicle (3A). Smoothing in-place also does not address gross anatomical misalignment. We address these issues in subsequent steps.

### Label refinement by detected anatomical edges

The extending, mirroring, and smoothing steps lead to a more complete set of labels for the ADMBA and correct the irregular borders in orthogonal planes to which the original labels were drawn; however, in several locations the labels do not align closely to the anatomical edges seen in the underlying histology images, for example the basal ganglia do not follow the curve of the lateral septal nuclei (Fig. 2C). To better map the anatomical fidelity of annotations in all dimensions, without manual relabeling, we leveraged our method for extending labels laterally based on gross anatomical edges to further refine all labels (Fig. 4).

**Figure 4:**
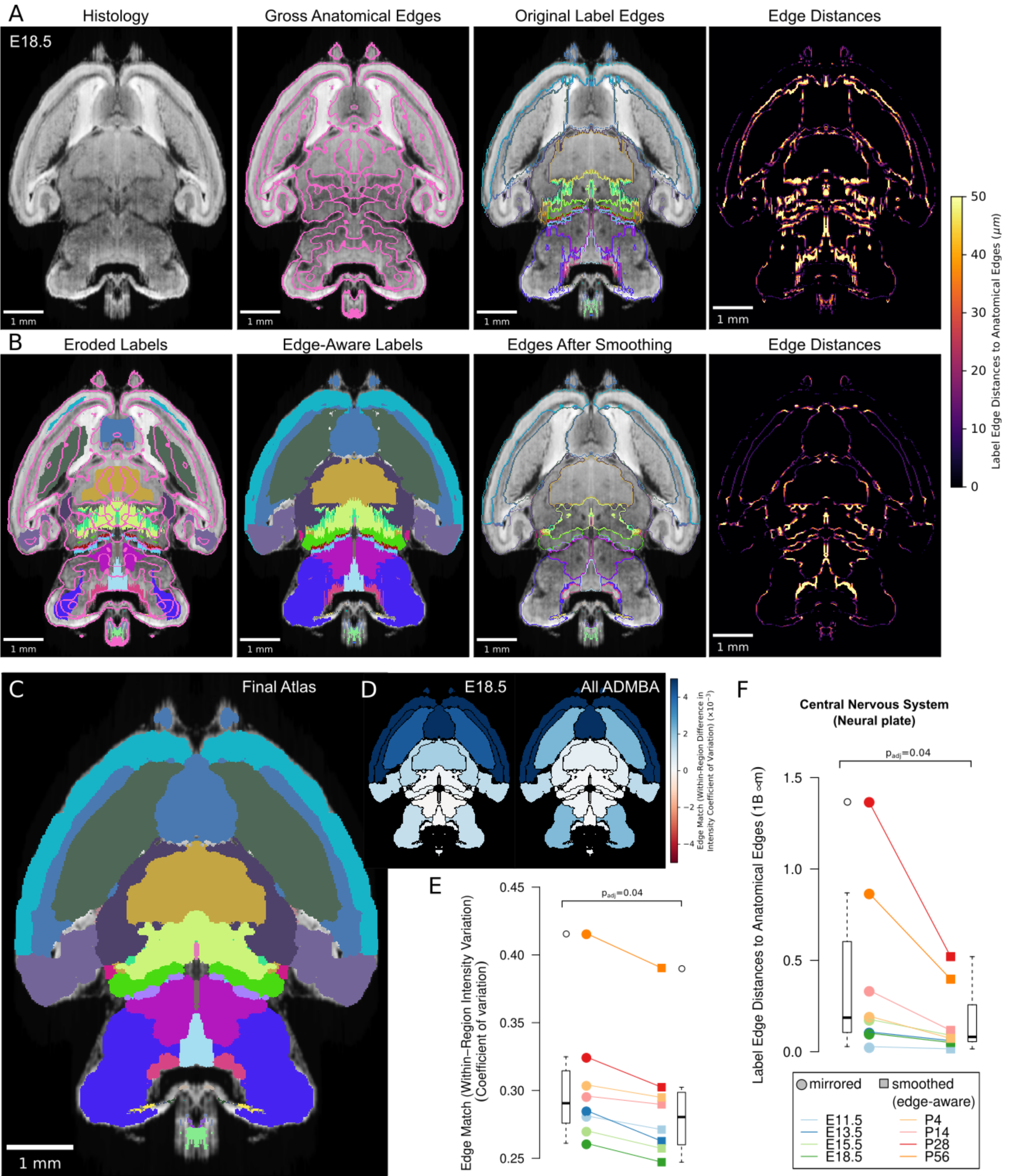
Edge-Aware Reannotation. (A) Edge detection of the volumetric histology image delineated gross anatomical edges (left), shown here with the E18.5 atlas. To compare these histology-derived anatomical edges with the extended and mirrored but unsmoothed label edges (center left), we used a distance transform method to find the distance from each label edge pixel to the nearest anatomical edge (center right), shown here as a heat map of edge distances (right). (B) Eroded labels served as seeds (left) from which to grow edge-aware labels through a watershed algorithm (center left), guided by the gross anatomical edges. After smoothing, borders matched the anatomical edges more closely (center right), as shown in the edge distance heat map for the modified labels (right), using the same intensity scale. (C) The final E18.5 atlas after edge-aware reannotation and label smoothing to minimize edge artifacts. (D) To evaluate the level of edge match by label, we mapped differences in the intensity coefficient of variation weighted by relative volume for each label before and after label refinement onto each corresponding label for the E18.5 atlas (left) and across all ADMBA atlases (right). For both, the anatomical map depicts this metric as a color gradient across all of the sublevel labels present in a cross section of the E18.5 atlas. Improvements of this metric with the refined atlas are colored in blue, minimal change is shown in white, while red represents better performance with the original atlas. (E) Applied across the full ADMBA, edge-aware reannotation and smoothing led to a significant improvement in the overall variation of intensities (central nervous system, or “neural plate,” level 0, ID 15565 in the Allen ontology; p = 0.04, n = 8 atlases, WSRT, Bonferroni corrected). (F) Distances from labels to anatomical edges similarly showed a significant improvement across atlases (p = 0.04, n = 8, WSRT, Bonferroni corrected).

Using the same gross anatomical map in 3D (shown for an example plane in Fig. 4A, second from left), we first quantified the distances from 3D label edges to the expected anatomical position. Assuming that the nearest gross anatomical edge was the correct one, we measured the distance from each label edge to the nearest gross anatomical edge. We can visualize this distance as a color gradient, in which higher intensity of color represents a greater distance to each anatomical edge (Fig. 4A, right columns).

To modify the labels in light of the gross anatomical edge map, we again incorporated the edge-aware algorithm, this time in 3D. We made the assumption that the core of each label is annotated accurately, whereas label edges are more prone to inaccuracies from plane-to-plane misalignments or the difficulty of assessing histological edges. To preserve the core while reannotating the periphery, we first eroded each label to remove edges. These eroded labels became the seed for the watershed step, which re-grew labels back toward their original size but now guided by the gross anatomical edge map.

Normally the erosion step would lead to loss of thin structures within labels because erosion operates globally on the entire label. To preserve these thin structures, we skeletonized each label in 3D, which thins the label to its core structure [58], and added the skeleton back to the eroded label. We used a much more lightly eroded version of the label for the skeletonization to avoid retaining label edges in the skeleton. By combining erosion with skeletonization when generating seeds for the watershed, we retained thin structures such as the alar plate of the evaginated telencephalic vesicle located anterior to the basal ganglia.

After performing this edge-aware step, we ran the adaptive morphological opening filter smoothing step (Fig. 4B). Because the edge-aware step partially smooths structures, we could use smaller filter sizes for smoothing to avoid loss of thin structures. The resulting labels show considerable improvement, for example the basal ganglia now curve around the lateral septal nuclei (Fig. 4C). Visualization of the color gradient of distances to anatomical edges also confirms substantial improvement in label alignment compared with the original labels or smoothing or edge-aware steps alone (Suppl. Fig. S8B). To quantify this improvement brain-wide, we calculated the sum of edge distances for each pixel at label surfaces across the ADMBA. We observed a significant reduction from a median of 186 million to 81 million *µm* (p = 0.04, WSRT, Bonferroni corrected; Fig. 4B, C, F), with a median Dice Similarity Coefficient between original (mirrored) and smoothed (edge-aware) labels of 0.72 (mean 0.77) and 9% median (mean 22%) volume reassignment. Example planes from all atlases in the ADMBA before and after refinement are depicted in Fig. 5, and movies across all planes are shown in Videos S1-16.

**Figure 5:**
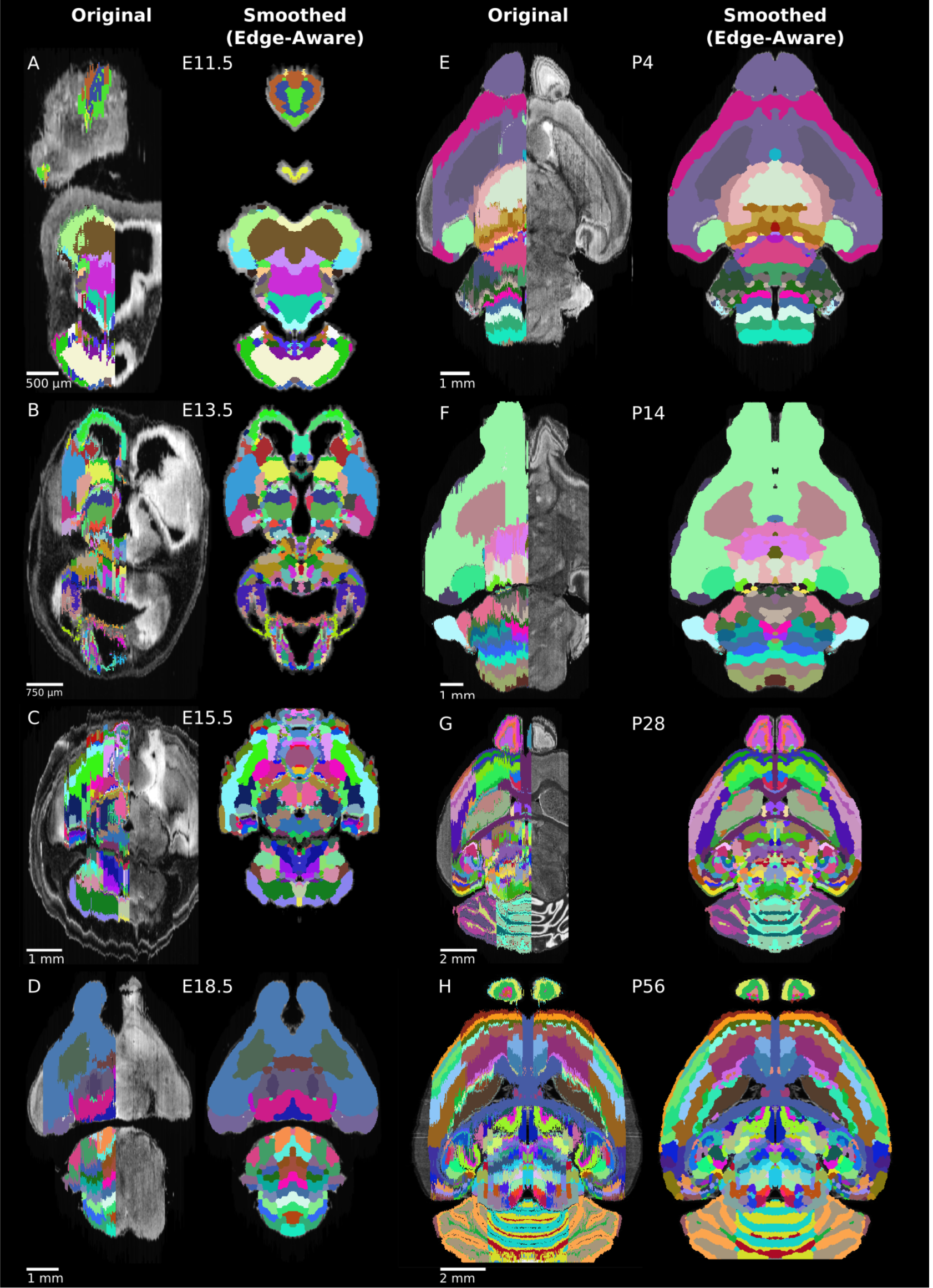
Refined Atlases. (a-h) Representative axial planes from all atlases across the ADMBA. For each pair of images, a plane of the original (left) atlas is depicted next to the refined (right) atlas after undergoing the full refinement pipeline. Complete atlases before and after refinement are shown as movies in Videos S1-16.

Anatomical edges in microscopy images reflect differences in intensity between regions. Therefore, we would expect accurate labels to have smaller variation in intensity in the underlying microscopy images than inaccurate labels, though such a difference would need to be apparent even though the majority of the label is unchanged. We used the coefficient of variation of intensity values within each label to quantify this expectation and demonstrated significantly lower variation with edge-aware labeling (mean from all labels weighted by size decreased from 0.305 to 0.289, p = 0.04, for all 8 atlases, WSRT, Bonferroni corrected; Fig. 4D, E). Furthermore, edge-aware labeling decreased the absolute coefficient of variation for 94 of the 100 individual labels represented by all atlases. The few labels that showed increased variation were frequently in regions of relatively subtle intensity changes, such as the hindbrain, where histological edges were less well-defined (Fig. 4D).

### Application to tissue-cleared whole brains

A major goal of 3D atlas refinement is for automated label propagation to optically sectioned, volumetric mouse brain microscopy images generated from intact tissue using recently developed tissue clearing techniques. As the accuracy of atlas registration is ultimately dependent on the fidelity of the underlying atlas in all dimensions, we sought to test and quantify improvement from our atlas refinements in cleared mouse whole-brains.

Among the many available methods to clear whole organs, we chose CUBIC [25], [59] given its balance of overall transparency, morphological retention with minimal tissue expansion, and ease of handling as a passive aqueous technique [60]–[62]. After clearing C57BL/6J WT mouse pup brains (age P0) with simultaneous incubation in SYTO-16 nuclear dye for 2 weeks, we imaged intact whole-brains by lightsheet microscopy at 5x to obtain volumetric images at cellular resolution, taking approximately 3h to image and generating approximately 500GB of data per brain (n = 15; 10 male, 5 female).

To detect nuclei throughout cleared whole-brains automatically, we implemented a 3D blob detection algorithm using the Laplacian of Gaussian filter, which has been shown to work well in equivalent image data

[63] (Fig. 6). To make the nuclei approximately spherical for blob detection, we interpolated the images axially to match the lateral resolution. Due to the large quantity of data, processing was performed in parallel on small chunks. Preprocessing and detection settings were optimized using hyperparameter tuning against a “truth set” of 1118 nuclei selected from multiple brain regions that had been verified by manual visualization (Suppl. Fig. S11A, B). The resulting model achieved a recall (sensitivity) of 90% and precision (positive predictive value) of 92%. Furthermore, the model showed high correlation with total intensity levels brain-wide (r *>* 0.99, p ≤ 1 × 10 ^−16^, for both original (mirrored) and smoothed (edge-aware) atlases) and nuclei vs. intensity densities (original: r = 0.89; smoothed: r = 0.93; p ≤ 1 × 10 ^−16^ for both), suggesting the performance was accurate outside the narrow target of the truth set (Suppl. Fig. S11C, D).

**Figure 6:**
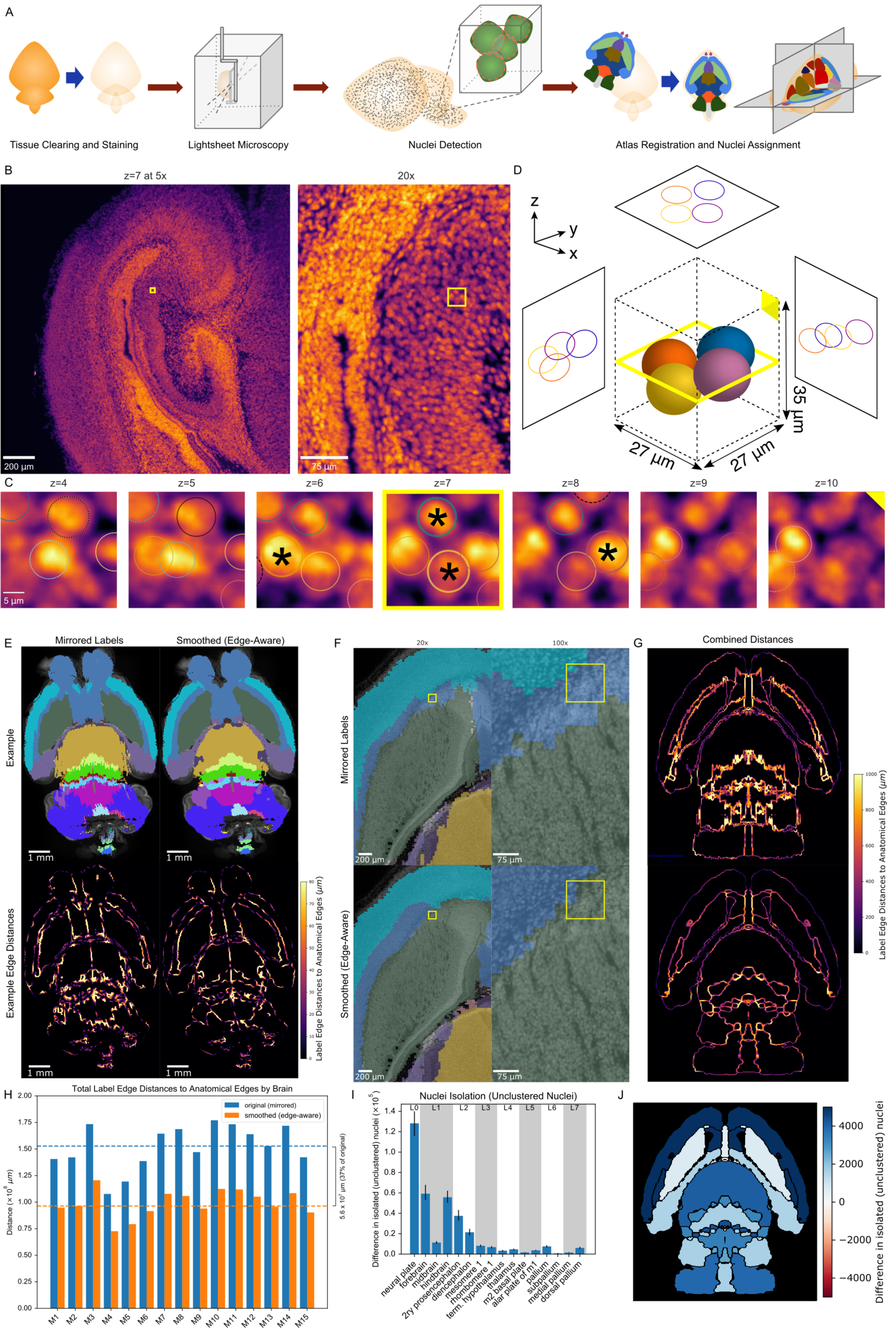
Nuclei Detection. (A) Overview of the nuclei detection and assignment pipeline. After tissue clearing by CUBIC and nuclear staining, we imaged intact whole P0 mouse brains by lightsheet microscopy. The E18.5 mouse brain atlas was registered to volumetric sample images. Nuclei were identified brain-wide with a 3D blob detector algorithm, and the nuclei coordinates were used to map to specific region labels, allowing quantification of nuclei counts and density for each label. (B) Axial view of the hippocampus of a P0 mouse brain at the original magnification of 5x (left) and zoomed to 20x (right), with the same region of interest (ROI) depicted by the yellow box in both. (C) Serial 2D planes along the ROI shown by the yellow box in part “B” show the emergence and disappearance of nuclei while progressing vertically (z-axis) through the ROI. Each of four individual nuclei is assigned a unique colored circle. The most central image of the nuclei is indicated by a solid circle and asterisk, while images above or below have dashed circles. (D) 3D schematic of the four nuclei from part “C” demonstrating their spatial orientation. (E) Example of the original (mirrored, left) and smoothed (edge-aware, right) E18.5 atlases registered to a representative cleared, optically-sectioned P0 wild-type mouse brain. Edge distances between the registered labels and the brain’s own anatomical edge map are reduced for this example brain, shown by the color gradient for each edge (bottom). (F) Label alignment at higher resolution. The top row depicts the registered original (mirrored) atlas at 20x and 100x around a region of interest highlighted by a yellow box. This same brain region is depicted in the bottom row, but overlaid with the refined atlas registered using identical transformations as in the original atlas. (G) Summation of the edge distances across all 15 wild-type brains with color gradient showing the edge distances with the original (top) and smoothed (bottom) labels. (H) Total edge distance at the whole-brain level before and after atlas refinement for each of these brains. (I) Density-based nuclei clustering within each label. Many of the isolated nuclei that could not be clustered in the original labels were clustered in the refined labels, with differences in unclustered nuclei shown from 16 regions selected across the hierarchy of labels from the grossest (neural plate, L0, left) to the finest (e.g. dorsal pallidum, L7, right). Error bars represent 95% confidence intervals. (J) The differences between the original and refined labels’ unclustered nuclei are depicted on an anatomical map showing this metric as a color gradient across all of the sublevel labels present in a cross section. Improvements with the refined atlas are colored in blue, while red represents better performance with the original atlas. A complete list of differences for each metric in each label is provided in Data S2.

Using the E18.5 brain as the closest atlas to our P0 C57BL/6J wild-type mouse brains (birth typically at E19 [64]), we then registered the atlas volumetric histology images to each sample brain. We chose to register brains using the Elastix toolkit [65], [66], which has been validated on CUBIC cleared brains and balances computational efficiency and accuracy [67]. After downsampling the volumetric images to the same dimensions as the atlas for efficiency, we applied rigid, followed by non-rigid, deformable registration based on a multi-resolution scheme, optimizing the cross-correlation similarity metric as implemented by the SimpleElastix [67], [68] programmable interface to the Elastix toolkit [65], [69]. After registration, the median DSC between the registered atlas volumetric histology images and the lightsheet microscopy images of each sample brain was 0.91 (mean: 0.91; 95% confidence interval: 0.90-0.92; Suppl. Fig. S10), although inspection of registered brains also revealed slight misalignments with the atlas, mostly in caudal regions including the cerebellum, a structure known to be challenging for registration [67]. Variations in sample preparation, including the completeness of the dissected hindbrain and expansion of the third ventricle during tissue clearing may also have contributed to these misalignments.

To evaluate whether our updated, smoothed 3D labels improved the analysis of true volumetric data, we registered both the original and refined labels to 15 tissue-cleared, nuclei-labelled whole wild-type mouse brains (10 male, 5 female). We would expect improved 3D labels to correspond more closely to the underlying anatomy. To test this expectation, we generated gross anatomical edge maps for each cleared brain and measured distances from label borders to anatomical edges. Edge distances significantly improved for almost all labels (overall decrease from a median of 153 million *µm* to 96 million *µm*, p = 0.007, WSRT, Bonferroni corrected for all hierarchical brain labels, mean 152 million to 99 million *µm*; Fig. 6E, G-H; Suppl. Fig. S13A). We would also expect improved 3D labels to have lower variation in image intensity within each label, as we observed in assessing the refined labels with the original brain images from which they were derived (Fig. 4E). We observed a small, but significant decrease in the intensity coefficient of variation at the whole-brain level (0.309 to 0.301, p = 0.007, WSRT, Bonferroni corrected, mean 0.311 to 0.304) and most sub-labels (Suppl. Fig. S13B). For 22 labels (18% of all labels) the variability worsened, however this was only significant (p = 0.04) for a single label. The majority of these 22 labels describe small structures (combined 7% of total volume) and therefore sensitive to slight perturbations in border location, and located ventrally in the brain, where signal was prone to being distorted by glue artifacts from the mount for lightsheet imaging. For context, variability improved for 98 labels (82% of all labels) with 27 labels showing significant improvement.

These two analyses support an overall improved performance of the refined 3D labels with the volumetric images, however we can also test nuclei density - a key use case for whole-brain imaging methods [33]. After detecting nuclei throughout each brain (Fig. 6), we assigned each nucleus to an atlas label based on the 3D location of the nucleus. Using the numbers of nuclei per label and the volume per registered label, we calculated nuclei densities using the original and refined labels (Suppl. Fig. S12). As with the volumetric image data, we would expect accurate labels to encapsulate regions with more constant nuclei density. We therefore assessed the coefficient of variation for nuclei density and observed a small, but significant improvement with the refined labels overall (median 0.629 to 0.625, p = 0.007, WSRT, Bonferroni corrected, mean 0.629 to 0.625) and across the majority of labels in the hierarchy (Suppl. Fig. S13C). A median of 13.5% (4.7 million) nuclei (mean 13.2% (4.6 million) nuclei) were reassigned from original to refined labels. Examples of these nuclei reassignments in a wild-type mouse forebrain is depicted in Fig. 6F. As an independent assessment of label alignment based on nuclei density alone, we measured nuclei clustering within each label under the expectation that well-aligned labels would group nuclei of similar densities, whereas poorly aligned labels would form isolated pockets of nuclei along borders between regions of contrasting nuclei densities. We employed Density-Based Spatial Clustering of Applications with Noise (DBSCAN) [70], an algorithm useful for both clustering and noise or anomaly detection [71], and indeed found that label refinement reduced isolated, unclustered nuclei by 30% (median 4.4 × 10^5^ to 3.1 × 10^5^ nuclei; p = 0.007, WSRT, Bonferroni corrected; mean 4.4 × 10^5^ to 3.1 × 10^5^; Fig. 6I-J), suggesting greater nuclei assignment to labels with nuclei clusters of similar density.

### MagellanMapper as a tool

To facilitate both automated atlas refinement and visual inspection of large 3D microscopy images, we developed the open-source MagellanMapper software suite. This Python-based [72], [73] image processing software consists of both a graphical user interface (GUI) for visualization and a command-line interface for automation (Suppl. Fig. S15). A major goal of the suite’s GUI is to enable unfiltered, raw visualization and annotation of original 2D images in a 3D context. The GUI consists of three main components: 1) A region of interest (ROI) selector and 3D visualizer with 3D rendering, 2) A serial 2D ROI visualizer and annotator, tailored for building truth sets of 3D nuclei positions, and 3) A simultaneous orthogonal plane viewer with atlas label editing tools, including painting designed for use with a pen and edge interpolation to smoothly fill in changes between two edited, non-adjacent planes. The command-line interface provides automated pipelines for processing of large (terabyte-scale) volumetric images, including 3D nuclei detection at full image resolution.

The suite makes extensive use of established computer vision libraries such as scikit-image [74], ITK [75] (via SimpleITK [76] and SimpleElastix [65], [68]), visualization toolkits such as Matplotlib [77] and Mayavi [78], and statistical and machine learning libraries such as Numpy [79], SciPy [80], Pandas [81], and scikitlearn [82]. The cross-platform capabilities of Python allow the suite to be available in Windows, MacOS, and Linux. We also leverage additional open-source imaging ecosystems such as the ImageJ/Fiji [83], [84] suite for image stitching through the BigStitcher plugin [85], integrated into our automated pipelines through Bash scripts.

## Discussion

3D reference atlases are critical resources for interpreting and comparing anatomically complex tissues, especially as techniques for whole-tissue microscopy [22], [33] and *in-situ* single-cell transcriptome methodologies are refined [86], [87]. Considerable effort has been invested in generating 2D reference atlases, while manually building atlases in 3D is time-consuming and laborious; building 3D atlases from these 2D resources will save time and improve comparability between analyses. Here, we have presented a method to automate the completion and conversion of atlases from 2D sections to complete, anatomically aligned 3D volumes and applied these methods to the eight developing mouse brain atlases in the ADMBA (Fig. 5).

Key features of our method include, first, an approach to adaptive morphology for label smoothing while preserving image detail by combining skeletonization, morphological erosion, and watershed segmentation. Second, we utilized this method as an edge-aware label refinement approach, which incorporates anatomical edge detection from an underlying intensity-based image to perform two functions: 1) Label extension into incompletely labeled regions, and 2) Anatomically guided label curation. Third, we integrated these approaches into an automated pipeline as a general way to refine serial 2D labeled datasets into 3D atlases. We next applied the pipeline to complete and refine the full suite of 8 atlases in the Allen Developing Mouse Brain Atlas series from ages E11.5 to P56 as a resource to the neuroscience and neurodevelopmental communities (Fig. 5). Specifically, these tools extended incomplete labels in each atlas to cover the entire brain and smooth artifacts between 2D sections, whilst respecting anatomical boundaries. We showed that the refined labels are more complete (Fig. 2), the interface between labels match anatomical images more closely (Fig. 4), and the image intensity within these labels is more homogeneous (Fig. 4). Using wholebrain lightsheet microscopy of 15 mice at P0, we identified the 3D location of 35 million nuclei per brain (median across 15 samples; Fig. 6). We showed that the refined labels match anatomical boundaries better at cellular resolution, providing a key step towards comparative analysis of whole brain cytoarchitecture (Fig. 6). Fifth, we have provided the method as an open source, graphical software suite with an end-to-end automated image processing pipeline for refining atlases, detecting nuclei, and quantifying nuclei counts by anatomical label in raw microscopy images of whole tissues.

Building a 3D reference atlas from a physically sectioned atlas presents three main challenges: 1) Incomplete labels resulting in unidentified tissues, 2) Smoothing jagged edge artifacts between sections in two of the three dimensions, and 3) Maintaining anatomical boundaries through the smoothing process. Prior work has tackled these challenges through two approaches. The first is automated smoothing of label boundaries [40], however this approach has well recognized limitations [88], including labels crossing anatomical boundaries, loss of label detail, and complete loss of small labels (Suppl. Fig. S4). The second approach uses a combination of novel data generation and copious, manual curation as applied by neuroanatomists and illustrators to version 3 of the Allen Common Coordinate Framework (CCFv3, 2017) [38], [47]. The resulting 3D reference atlas is highly accurate; however, an even greater large-scale effort would be required for the eight stages of development in the ADMBA.

Our method automates additional curation steps to extend and smooth labels guided by the underlying anatomy providing an automated method closer to the accuracy of manual curation. To address loss of detail and loss of labels, we used the combination of adaptive morphology and an edge-aware watershed to preserve small structures during automated smoothing. By fitting and extending labels to anatomical boundaries, we improved both the accuracy and label coverage of the resulting 3D reference atlas, without requiring additional data. While manual curation remains the gold standard as performed in the Allen CCFv3, even this monumental effort translated only 77% of labels because of resource limitations [47]. As suggested by Wang et al., future atlas development lies in automated, unbiased approaches [47]. We demonstrate qualitatively (Fig. 5) and quantitatively (Fig. 4) that our automated approach is a substantial refinement over existing reference atlases and creates a complete resource that is immediately applicable for registration of 3D imaging data (Fig. 6). Furthermore, it acts as a foundation for further improvements, substantially reducing the work required to generate a manually curated atlas in the future while freeing up anatomists and neuroscientists for the role of oversight in such endeavors.

Despite these substantial improvements, we are aware of some specific limitations of the approach. We expected less variability in image intensity within a region than between regions, and therefore improved region labels should reduce image intensity variability. While this expectation was correct for the majority of labels (82%), a small number of regions showed a modest increase in variability with our refined labels. Most of these labels were in the hindbrain, where histological contrast is lower, leading to less well-defined anatomical edges. Adjusting the weight of edge-detected boundaries based on the degree of underlying intensity contrast could lessen the impact of these edges on reannotation. A few more labels exhibited this paradox when registered to the wild-type brains, likely owing to artifacts from tissue preparation and mounting, the slight age mismatch between the closest available E18.5 atlas and the P0 wild-type brains, registration misalignments, and inaccuracies of the blob detector.

Another limitation is the use of a single filtering kernel size for smoothing across the entire brain (Fig. 3). In labels with both thick and thin portions, the thin portions are prone to being lost during the erosion step of smoothing. While a fully adaptive morphological approach would be beneficial, our use of skeletonization and the edge-aware watershed allowed us to preserve the core of such thin portions. More generally, the combination of erosion, skeletonization, and watershed could serve as an adaptive form of morphological operations to better preserve any label’s overall footprint and topological consistency.

While the refined labels are superior to the original ones in both the ADMBA images and whole-brain lightsheet microscopy images, we observed more registration artifacts with the whole-brain lightsheet microscopy images (Fig. 6E). Undoubtedly, biological variation, the slight age mismatch (E18.5 in ADMBA vs. P0 for lightsheet), and the different preparations and imaging modality (Nissl sections vs. nuclei-stained whole-brain) are substantial factors. Generating a group-wise composite of nuclei-stained whole-brain could help reduce biological variation. Deferring the edge-aware reannotation step until after registration using boundaries identified in the nuclei-stained whole-brain may also reduce these artifacts.

Recent methodological advances may further improve the refined 3D labels. Chon et al. integrated the Franklin-Paxinos atlas with the CCFv3 atlas and additional cell-type specific markers, addressing discrepancies between these standard references [2]. Integrating our anatomical edge maps with *in-situ* hybridization expression patterns [38], [46] may similarly improve label boundaries. A deep learning model utilized histological textures to generate labels [88]. Training the model required substantial manual annotation to serve as the initial ground truth; in this context, our automated approach to atlases generation may be able to replace or augment this manual step. Another recent approach used fractional differential operators to preserve texture as well as edge information for segmentation of single-cell resolution data [89]. This could be incorporated into the 3D anatomical edge map generation step to further delineate labels at higher or even full resolution.

## Conclusion

Mouse whole-brain imaging has been used to understand models of human traits, including sexual dimorphism [7] and models of human disorders, including Alzheimer’s disease [90]–[92], serotonin dysregulation [93], epilepsy [94], and Autism Spectrum Disorder (ASD) [95]. Such analyses would be augmented by accurate 3D reference atlases allowing the detection of subtle quantitative changes not readily appreciable in individual slices. As we have described and demonstrated here, anatomically guided completion and refinement of reference atlases more fully leverages the many existing and established atlases by expanding them to full coverage and greater accuracy at cellular resolution. The completed and refined 3D ADMBA serves as a resource to help identify biological differences in the multiple models of human disorders and traits across brain development.

## General

We also grateful for the microscopy advice and expertise from Meredith Calvert, director of the Gladstone Institutes Histology and Light Microscopy Core. We greatly appreciate Eric Lam from the KAVLI-PBBR Fabrication and Design Center at UCSF for assistance with the initial design of the custom 3D printed microscopy brain mount. We are deeply indebted to Hwee Kuan Lee from the Agency for Science, Technology and Research (A*STAR) for invaluable imaging informatics advice. We appreciate support from the Institute of Molecular and Cell Biology at A*STAR.

## Author contributions

DMY developed the refinement methods, performed imaging, and co-wrote the associated manuscripts. SFD performed mouse husbandry and tissue clearing and assisted with imaging. GS assisted with tissue clearing and imaging. ZB piloted the tissue clearing and imaging. DY and MN built nuclei ground truth sets and tested the software. LG provided advice on image processing methods. JR provided expertise on mouse development and anatomy. WY provided guidance on the method development. SJS conceived of the experiment, obtained funding, provided strategic guidance on the entire project and co-wrote the associated manuscripts.

## Data Availability

The method is freely available as open source software at: https://github.com/sanderslab/magellanmapper. Note to editors and reviewers: The refined ADMBA atlas files are too large to be included as supplements. We will upload the full atlases to the Human Brain Project EBRAINS data platform (https://ebrains.eu/) [96] and include the accession numbers in the publication. Videos comparing the original and 3D reconstructed atlases are available as supplementary files and here: https://sanderslab.github.io/data/.

