## Supplemental methods and figures for "Constructing and Optimizing 3D Atlases From 2D Data With Application to the Developing Mouse Brain"

### Materials and Methods

#### Original atlases

The ADMBA series provides atlases for multiple developmental time points from embryonic through adult stages [46]. Each atlas consists of two main images given in a volumetric format (.mhd and its associate .raw file), a microscopy image ("atlasVolume") and a labels image ("annotation"), which are each composed of multiple 2D sagittal planes. As outlined in the ADMBA technical white paper [97] and online application programming interface (API) documentation [98], each microscopy plane is from imaging of a C57BL/6J mouse brain specimen cryosectioned into 20 $\mu m$ (E11.5-P4) or 25 $\mu m$ (P14-P56) thick sagittal sections stained with Feulgen-HP yellow nuclear stain (E11.5-E18.5) or Nissl staining (P4-P56). The annotation planes correspond to each microscopy plane, with integer values at each pixel denoting the label at the finest structural level annotated for the corresponding microscopy voxel. To create the atlas labeling, an expert anatomist annotated 2D planes with Adobe Illustrator CS [50].

#### Lateral labels extension

While the ADMBA covers a large proportion of unique areas within each brain, the atlas labels leave the lateral edges of one hemisphere and the entire other hemisphere unlabeled. To fill in missing areas without requiring further manual annotation, we made use of the existing labels to sequentially extend them into the lateral unlabeled histology sections. This process involves 1) resizing the last labeled plane to the next, unlabeled plane, 2) refitting the labels to the underlying histology, and 3) recursively labeling the next plane until all unlabeled planes are complete (Suppl. Fig. S1).

To extend lateral edges of the labeled hemisphere, we first identified the lateral-most labeled plane of the atlas microscopy images. We started from the first sagittal plane on the labeled side of the image and moved toward and into the brain, checking each plane for the presence of any label. Once we identified a contiguous stretch of planes with labels, we used the most lateral labeled plane as the template for subsequent labels to be extended out laterally, in the opposite direction. In a few atlases (e.g. P28), the lateral-most labeled planes are only partially complete, in which case the most lateral completely labeled plane was manually specified instead.

This last labeled lateral plane contained one or more discrete structures to extend label coverage, typically the cortex and sometimes the cerebellum. To find each structure and its associated labels, we first generated a mask of the histology by taking a slightly dilated version of the labels (`morphology.binary_dilation` method in scikit-image), typically with a disk shaped structuring element of size 5) to capture nearby unlabeled histological structures, thresholded this histology plane by a value of 10, removed small objects of size less than 200 connected pixels (default connectivity of 1 in `morphology.remove_small_objects`), and identified bounding boxes of connected histology components (`measure.regionprops`) and their matching labels. These bounding boxes contain the discrete structures that we will follow laterally, using the corresponding labels for each structure as templates to extend into the rest of each structure.

After identifying each structure and its labels, we fit the labels to the next lateral plane and recursively generated a new template for the subsequent plane. To fit the labels to the next plane, we found the histology bounding box for each structure and resized its labels to this box. We assumed for simplification that each structure is the same or smaller size in each subsequent plane, such as the tapering profile of the cortex laterally, and took only the single largest object found within the structure's bounding box. Spline interpolation, anti-aliasing, and range rescaling were turned off during the resize operation to avoid introducing new label values. Some atlas labels contain empty space such as ventricles. To ensure that the ventricles close as they progress laterally, we employed an in-painting approach. Using the Euclidean distance transform method from the Scipy [80] library (`ndimage.distance_transform_edt`), we identified the indices of the nearest neighboring pixel for any unlabeled pixel whose corresponding histology intensity value was above threshold and filled this missing pixel label with the value of this neighbor.

### Edge map generation

While the labels from one plane could serve as an approximate template for the next plane, we curated this template to fit the underlying anatomy. To map this anatomy, we generated gross anatomical edge maps of the histology images through 3D edge detection. First, we smoothed the volumetric microscopy image using a Gaussian filter with a sigma of 5 followed by an edge detection with a Laplacian filter using a default operator size of 3, using both filters implemented in scikit-image (`filters.gaussian` and `filters.laplace`, respectively). This relatively large Gaussian sigma allowed for capture of broad anatomical edges such as the cortex and basal ganglia. To enhance edge detection of the outermost boundaries, we identified and removed background by thresholding the original microscopy image with an Otsu threshold (`filters.threshold_otsu`) and combining it with a mask of all the original labels to fill in any missing holes in the thresholded image.

Finally, we reduced the edge-detected image to a binary image of edges alone by applying a zero-crossing detector. This detector separately erodes and dilates (`morphology.erosion` and `morphology.dilation`, respectively, with a ball-shaped structuring element of size 1) the edge-detected image to find borders, taking all pixels where this image changed signs as gross anatomical edges (Fig. 4A).

### Anatomically guided serial 2D reannotation

This edge map allowed us to conform labels to local anatomy. To refit labels, we eroded and regrew them in a watershed transformation guided by the anatomical edge map.

First we eroded labels individually (`morphology.binary_erosion` in scikit-image with a ball-shaped structuring element of manually determined radii for each atlas). These eroded labels served as seeds for a compact watershed (`morphology.watershed` implemented in scikit-image with a `compactness` parameter of 0.005) [54], [99], guided by the anatomical edge map. We used the Euclidean distance transform of the anatomical edge map to define the catchment basins for the watershed transformation, where voxels farther from anatomical edges are at the bottoms of basins and fill faster, guiding the regrowth of eroded labels. Labels typically crossed several anatomical edges but tended to meet neighboring labels at common edges (Fig. 4B). To limit the watershed to atlas foreground, which prevents label spillover across empty spaces, we set the watershed `mask` parameter to the original total labels foreground smoothed by an opening filter (`morphology.binary_opening` with a ball structuring element of radius 2) to remove artifacts around small ventricular spaces that might otherwise be crossed. The eroded labels thus regrew to fit anatomical guides and became the new, anatomically-refined template to extend labels into the next plane.

To model the tapering and disappearance of labels laterally, we allowed labels to erode completely. We weighted erosion toward central labels within each structure by multiplying the erosion filter size for each label by the label's median distance to the structure perimeter divided by the maximum distance. Instead of simply eroding away the smallest labels, this approach preferentially eroded labels farthest from the perimeter, typically central labels.

In atlases such as the ADMBA E18.5 atlas, a few spurious labeled pixels from other regions regrew to create artifacts. To filter these spurious pixels, we applied smoothing to the initial labels template using the smoothing approach described below. Also, we applied skeletonization as outlined below to avoid loss of thin structures during erosion.

Each plane of labels thus conformed to its underlying anatomy and became the template for the next plane, keeping labels inherently connected from one plane to the next. While this approach is in serial 2D rather than fully 3D because the starting labeled plan is 2D, a subsequent step will further refine labels in 3D.

### Atlas 3D rotation and mirroring

To fill the missing hemisphere in the labels image, we initially simply mirrored the present labels and underlying histology planes across the first unlabeled sagittal plane on the opposite side. We noticed however that this mirroring frequently duplicated midline structures because the labels extend slightly past the true sagittal midline. When we shifted the mirroring to the sagittal plane closest to midline, we found that many labels were lost in the final image.

Many of the developing atlases contain at least a few labels positioned solely across the midline in the otherwise unlabeled hemisphere. While it is possible that these labels represent structures unique to one hemisphere, at least some of these labels are on the other side of the midline in other atlases of the ADMBA and thus more likely represent artifact. In some cases (e.g. P4 and P14), mirroring just slightly past the sagittal midline preserved these labels, whereas other atlases (e.g. P28) contained over 100 labels past the midline. One approach to preserve these labels would be to compress the near-midline labels to bring all labels into a single hemisphere at the expense of potentially misaligning otherwise appropriately positioned labels. To avoid this side effect, we elected to cut off a few labels to preserve the placement of the majority of labels.

Upon closer inspection, many of the brains are slightly rotated in 2 or 3 dimensions, which contributed to loss of midline labels during mirroring. To rotate images volumetrically in 3D, we applied the scikit-image `rotate` function to all 2D planes along any given axis for each rotation. For each atlas, we applied this volumetric rotation along all necessary axes until the sagittal midline was parallel to an image edge, manually inspecting each brain in our 2D/3D orthogonal viewer to ensure symmetry. In some cases, such as the E18.5 atlas, the brain skewed laterally along its rostrocaudal axis. To avoid introducing a gap between midline labels and the corrected midline after rotation, we filled all planes on the unlabeled hemisphere side with the last labeled plane before rotation. Mirroring would then overwrite all of these repeated labels except those that filled in potential gaps left by rotation. After rotation, we noticed that mirroring reduced label loss in at least some atlases (e.g. E13.5).

### Piecewise 3D affine transformation

The distal spinal cord of the E11.5 atlas is strongly skewed and would be duplicated during mirroring. The skew is complicated by the cord's coiling back on itself and progressive skew along its length. To bend the distal cord to the sagittal midline, we developed a method for piecewise 3D affine transformation. This transformation allows a cuboid ROI within a volumetric image to be sheared while maintaining a specified attachment point along another axis, reducing discontinuity with neighboring areas.

First, we specify an axis along which to shear and the degree of shearing for a given ROI. Each full plane along this axis is shifted in the indicated direction by a progressively larger amount, overwriting pixels into which the plane is shifted and filling vacated pixels with background to shear the stack smoothly in 3D. If an axis of attachment is also specified, each plane is sheared line-by-line along this axis so that the resulting parallelogram remains fully connected at one end of each axis. The resulting ROI is thus sheared in 3D while remaining smoothly connected to its surrounding space along two orthogonal faces of the original cuboid ROI to minimize disruption.

Applied to the skewed spinal cord, this approach allowed us to shear the cord one section at a time as it curved back along itself (Suppl. Fig. S6, left column), with the connected faces ensuring that each piece remains attached to one another. First we sheared the entire distal cord from the start of the skew (Suppl. Fig. S6, middle left column). This shift brought the proximal section toward midline, but the more distal cord remained skewed where it curved back on itself. To correct this skew, we sheared again but starting from a more distal point and along an orthogonal angle to follow the cord's curve (Suppl. Fig. S6, middle right column). Now most of the cord lined up with the sagittal midline, but the shear exacerbated the skew of the most distal cord. Finally, we applied a third affine, this time on only the most distal section and in the opposite direction as the prior affine (Suppl. Fig. S6, right column).

After mirroring the resulting brain and cord, no duplication could be seen. This piecewise, overlapping affine of targeted regions allowed straightening of the cord without breakages or alteration of surrounding areas. Although surrounding non-CNS tissue suffered noticeable breakage, they were stripped out as described below.

### Stripping non-CNS signal

The extended labels allowed us to mask and crop out non-CNS tissue, including the rest of the embryo present in several of the embryonic stage atlases. While distinguishing this tissue based on intensity characteristics alone would be challenging, especially for areas where the spinal cord extends the length of the embryo, the extended, mirrored labels provide a map of relevant CNS tissue.

For each of the atlases with non-CNS tissue (E11.5-E15.5), we first cropped the atlas to the bounding box of these labels along with a small padding (5px) to remove much of the non-CNS tissue, such as the entire unlabeled body in the E15.5 atlas. We removed non-CNS tissue remaining within the cropped areas by using the labels as a mask to remove all histology pixels outside of the labels mask, including the body surrounding the labeled spinal cord in E11.5. To avoid missing tissue that may have been unlabeled, we dilated the labels mask slightly (`morphology.binary_dilation` with a ball-shaped structuring element of size 2) so that the mask encompasses both the labels and its immediate surroundings. The resulting histology thus contains all pixels in the near vicinity of labels, including pixels that should be labeled but are not.

### Label smoothing

While labels appear generally smooth when viewed from the sagittal plane, label edges are noticeably jagged when seen from the orthogonal directions. A previous smoothing solution proposed by Niedworok et al. [40] utilized a Gaussian blur with a sigma of 0.5 in two iterations to minimize ragged edges in an atlas derived from the Allen Reference Atlas P56 mouse brain atlas. We applied this approach by iteratively applying the Gaussian filter implemented in scikit-image (`filters.gaussian`) with a range of sigmas (0.25 to 1.25) to each label in 3D (Suppl. Fig. S4).

To extract each label in 3D, we found the bounding box of the label using the scikit-image `measure.regionprops` method and added additional empty padding space around the box. The Gaussian filter was applied to the label, and the original label in the bounding box was replaced with the smoothed label. Each label smoothing left small gaps vacated by the previously ragged borders. To fill in these gaps, we employed the in-painting approach described above. Finally, we replaced the original bounding box with that of the smoothed label in the volumetric labels image. We repeated the process for all labels from largest to smallest to complete the smoothing.

To enhance smoothing while retaining the original underlying contour of each label, we devised an adaptive opening morphological filter approach. The morphological opening filter first erodes the label to remove artifacts such as ragged edges, followed by dilation of the smoothed label to bring it back toward its original size. In place of the Gaussian filter, we applied this opening filter (`morphology.binary_opening` in scikit-image). For smaller labels, the filter occasionally caused the label the disappear, particularly for sparsely populated labels with disconnected pixels. To avoid label loss, we employed an adaptive filter approach by halving the size of the filter's structuring element for small labels ($\leq$5000 pixels). For any label lost in spite of this filter size reduction, we switched the filter to a closing morphological filter (`morphology.binary_closing`), which dilates first before eroding and tends to preserve these sparse labels at the expense of potentially amplifying artifact.

### Smoothing quality metric

To evaluate the quality of smoothing and to optimize morphological filter structuring element sizes, we developed a smoothing quality metric with the goal of balancing smoothness while maintaining the overall original shape and placement. Since the major effect of label smoothing is to minimize high frequency aberrations at label borders and thus make each label more compact, we measured this amount of smoothing by the 3D compactness metric. We termed the difference in compactness before and after smoothing as "compaction." As the goal of smoothing is to remove these artifacts while preserving the overall shape of the label, we introduced a penalty term of "displacement," measured by the volume shifted outside of the label's original bounds.

We used the classical measure of volumetric compactness, where lower values are more compact [55]:

$$\text{Compactness} = \text{SA}^3/\text{Vol}^2 \tag{1}$$

To measure the surface area of each label in 3D, we employed the marching cubes algorithm [100] as implemented in scikit-image (`measure.marching_cubes_lewiner`), which also accounts for anisotropy. The algorithm extracts surfaces in 3D by dividing the volume into cubes and marching through these cubes to find where isosurfaces intersect with each cube, forming triangular surfaces within each cube wherever an isosurface passes through the cube. Using a mask of each label, we obtained the edge surface as a mesh from which we measured the surface area (`measure.mesh_surface_area`). We took the volume as the total number of mask pixels and multiplied by the product of the voxel spacing to account for anisotropy. With the surface area and volume, we calculated the labels compactness (1). To quantify the fractional change in compactness before and after smoothing, we took the original compactness minus the smoothed compactness and normalized the difference to the original compactness to give the unitless value that we termed "compaction":

$$\text{Compaction} = \frac{\mathrm{C}_{\mathrm{orig}} - \mathrm{C}_{\mathrm{smooth}}}{\mathrm{C}_{\mathrm{orig}}} \tag{2}$$

where $\mathrm{C}$ is the 3D compactness given above.

To measure "displacement," we measured the volume shifted outside of the label's original bounds. Taking the mask of the smoothed label, we combined it with the inverse of the mask of the original label through a boolean `and` before totaling the number of pixels. Similarly to the compactness measure, we normalized the displacement volume to the original volume:

$$\text{Displacement} = \frac{\mathrm{Vol}_{\mathrm{smooth}} \notin \mathrm{Vol}_{\mathrm{orig}}}{\mathrm{Vol}_{\mathrm{orig}}} \tag{3}$$

where $\mathrm{Vol}$ is the volume of the given label.

As a measure of smoothness quality, we took the difference of the compaction and displacement:

$$\text{Smoothing quality} = \text{Compaction} - \text{Displacement} \tag{4}$$

To quantify the smoothing quality for the entire atlas, we took the weighted arithmetic mean of all the labels' smoothing qualities, weighting by each label's volume. The atlas-wide smoothing quality metric can be summarized in the following equation:

$$\text{Atlas-Wide Smoothing Quality} = \frac{\sum_{i=1}^{N} \mathrm{SmoothingQuality_i} \mathrm{Vol_{i,orig}}}{\sum_{i=1}^{N} \mathrm{Vol_{i,orig}}} \tag{5}$$

where N is the total number of original labels. While maximizing compaction would reduce surface area the most by transforming the label into a perfect sphere, the displacement from the label's original space would penalize this over-compaction, allowing us to target the balance of compaction and displacement to find the optimal smoothing quality.

### Anatomical to label edge distance quantification

As an automated method of quantifying the correspondence between labels and anatomical edges, we used the anatomical edge maps generated earlier to measure the distance between anatomical and label edges. We reduced each label to its edges in 3D by eroding the very outer surface of each label (`morphology.binary_erosior` with a cross-shaped structuring element with connectivity of one) and subtracting this eroded label from the original label.

To measure distances between label and anatomical borders, we performed the Euclidean distance transform (`ndimage.distance_transform_edt` in Scipy) on the anatomical edge map to measure distances from any given voxel to the anatomical edges, using the `sampling` parameter to specify the microscopy image spacing in $\mu m$. Using the labels edge map, we next took only the voxels in the distance map corresponding to these label borders as a map of distances from each label voxel to its nearest anatomical edge. As overall measures of distance from label borders to the nearest anatomical border, we summed the edge distances for each label to compare before and after label refinement.

### Anatomically guided 3D reannotation

Generating edge images provided a map of the boundaries between anatomically distinct regions not only to measure distances between borders, but also to curate the labels themselves with these anatomical edges. To reannotate the existing unsmoothed labels, we eroded and regrew them through an anatomically guided watershed transformation similar to the approach described above but now in 3D. We tested seeds with multiple erosion filter sizes and found that a structuring element size of 8 reduced the seed size sufficiently to correct label bounds around several grossly abnormal labels, including the lateral septal nuclei and basal ganglia.

As a global operator, erosion typically leads to loss of thin structures, especially with larger structuring elements. To avoid this loss, we first extracted the core structure of each label by finding its 3D skeleton (`morphology.skeletonize_3d` in scikit-image). We added the skeletonized image back to the eroded label to recover the location of thin structures, allowing the watershed to regrow these labeled areas in addition to the eroded label. To limit branches in the skeleton, which could counter the effect of the erosion, we input a lightly eroded version of the labels (structuring element half the size of that of the main erosion) to the skeletonization.

The resulting watershed segmentation also served as preliminary smoothing but introduced its own label edge artifacts, though of lower frequency than in the original labels. We thus deferred the smoothing algorithm until after this watershed step and could use smaller filter sizes to generate a final smooth image.

### Application to the full ADMBA series

Each atlas in the ADMBA required a different set of refinement features and settings, including specialized adjustments such as the 3D piecewise affine only for a specific atlas (E11.5). To allow for customized settings, we defined separate profiles of parameters within our software suite for each atlas.

### E11.5

The microscopy images depict a specimen in embryonic stage with the caudal end including the spinal cord wrapping around itself and deviated laterally from the rest of the body. As with most other atlases in this series, one half of the labels were missing. Making the image symmetric would allow us to mirror labels from the existing side onto this missing side and minimize bias when using the atlas for automated registration tasks. We started by rotating the image by 5° in the axial planes toward the left of the brain and 1° in the coronal planes to raise the right side of the brain, bringing the sagittal midline parallel to an image edge. To shift the distal end of the spinal cord back toward the sagittal midline, we applied the piecewise 3D affine transformation described above (Suppl. Fig. S6). Prior to the affine, we applied an additional rotation of 30° in the sagittal planes to position the skewed distal cord within a single cuboid ROI parallel to the image.

With the embryo now symmetric, we mirrored the labels and microscopy signal along the embyro's midline, measured as 52% across the sample along the z-axis. To highlight the central nervous system and remove breakages introduced by the affine transformations in non-CNS tissue, we stripped out non-CNS tissue as described earlier.

The E11.5 atlas uniquely contains labeled ventricles. To ensure that the Laplacian of Gaussian edge-detection algorithm appropriately found ventricular edges, we used only the thresholded atlas rather than incorporating the labels to find the background. We also included the ventricular space as foreground for purposes of the DSC calculations between microscopy and labels to account for the ventricular labeling. This atlas did not require lateral extension.

### E13.5

This atlas also contains the full embryo, but the spinal cord is symmetric along the sagittal midline of the brain and did not require the affine transformations as in the E11.5 atlas. After extending the lateral edges, we rotated the atlas by 4° in the axial planes toward the left of the brain and 2° in the coronal planes to lift the right side of the brain, making the atlas symmetric before mirroring the atlas at 48% along the sagittal planes. We again stripped non-CNS tissue the same way as for the E11.5 atlas.

### E15.5

This atlas is the final one to include the complete embryo, although only the very rostral end of the spinal cord includes labels. We extended the lateral edges, rotated the images by 4° in the axial planes toward the left side of the brain, mirrored the atlas at 49% along the sagittal planes, and stripped away the entire embryo outside of the brain. A stepwise shift in sagittal planes is apparent at several planes (e.g. 104 and 129) in both the histology and label images in the original atlas, which we smoothed slightly in the labels during the smoothing step.

### E18.5

A small subset of labels from sagittal planes 103-107 were compressed along the dorsoventral axis. To match them with their neighboring label planes and the underlying atlas, we resized them similarly to the lateral edge extension. Starting with the first plane in this subset, we thresholded the microscopy image, removed small objects, and obtained the largest bounding box of connected histology components. Taking only this largest connected structure allowed us to avoid including extraneous tissue visible on the ventral aspect of the brain, which was unlabeled. We repeated the process on the labels to obtain its compressed bounding box and resized it to the size of the microscopy bounding box. Finally, we repeated the entire process on the rest of the planes in this subset (S7).

For the lateral edge extension, we noted that the basal ganglia in the most lateral planes are slightly larger than in more medial planes. Under the assumption that the basal ganglia would be tapering laterally, we skipped these planes and started the extension at 13.7% along the sagittal planes to take the plane with smallest basal ganglia label. After rotating the atlas by 1.5° in the axial planes toward the right of the brain and 2° in the coronal planes to lift the left side of the brain, we mirrored microscopy and labels at 52.5% along the sagittal planes for symmetry.

**P4**

This atlas is reminiscent of E18.5 but without the necessity of setting the lateral edge extension starting plane explicitly or expanding any compressed labels. After lateral edge extension and rotating the atlas by 0.22° in the axial planes toward the right of the brain, we mirrored microscopy and labels at 48.7% along the sagittal planes for symmetry.

**P14**

The most laterally labeled plane is discontinuous with the rest of the labeled planes in this atlas, so we skipped this plane during lateral extension and started extending only from the first contiguous set of planes. After rotating the atlas by 0.4° in the axial planes toward the left of the brain, we mirrored the microscopy and labels images at the 50% mark along the sagittal planes.

**P28**

The most lateral planes had incomplete labels, requiring use of a more medial plane with complete labels at 11% along the sagittal planes for the lateral edge extension. After rotating the atlas by 1° in the axial planes toward the right of the brain, we mirrored the microscopy and labels images at the 48% mark along the sagittal planes.

## P56

The ADMBA contains a P56 mouse similar to the adult P56 but following the same ontological labeling scheme as in the rest of the ADMBA. This atlas uniquely contains bilateral labels, although the far lateral section is still missing. We again extended the lateral edges and mirrored the microscopy and labels images, starting extension at 13.8% and mirroring at 50% along the sagittal planes, respectively. The most lateral labeled plane contains two distinct labeled structures, the cortex and cerebellum, requiring separate extension for each distinct structure as outlined in our method above.

### Animals and tissue clearing

All procedures and animal care were approved and performed in accordance with institutional guidelines from the University of California San Francisco Laboratory Animal Research Center (LARC). All strains were maintained on a C57BL/6J background. Animals were housed in a vivarium with a 12h light, 12h dark cycle. For timed pregnancies, noon on the day of the vaginal plug was counted as embryonic day 0.5. Pups were harvested at P0 (postnatal day 0).

At the time of experiment, P0 pups were anesthetized on ice and perfused transcardially with ice-cold 1X PBS supplemented with 10 U/mL heparin and then with 4% PFA in 1X PBS, followed by brain isolation. P0 brains were post-fixed overnight at 4°C in 4% PFA in 1X PBS. The next day the excess fixative was removed by washing the brains with 1X PBS supplemented with 0.01% (wt/vol) sodium azide (Sigma-Aldrich) for at least 2h at room temperature (RT).

Samples were cleared using the advanced CUBIC clearing protocol for whole-brain and whole-body clearing [59]. In short, samples were immersed in 1/2-water-diluted Reagent-1 containing 1 $\mu M$ SYTO-16 (Thermo Fisher) and incubated at 37°C for 6h (Reagent-1: 25 weight% (w%) Urea, 25 wt% Quadrol, 15 wt% Triton X-100, and dH2O). 1/2 diluted Reagent-1 was replaced with Reagent-1 containing 1 $\mu M$ SYTO-16 and incubated overnight at 37°C on a rotator. The next day, solution was replaced with fresh Reagent-1 containing 1 $\mu M$ SYTO-16 and was incubated overnight at 37°C. Reagent-1 was replaced with fresh Reagent-1 containing 1 $\mu M$ SYTO-16 every 48 hrs for a total of 8 days. Tissue clearing was stopped by washing the sample with 1X PBS supplemented with 0.01% (wt/vol) sodium azide at RT once for 2h, once overnight and again once for 2h. After the wash step, samples were immersed in 10 mL of 1/2-PBS-diluted reagent-2 (Reagent-2: 25 wt% Urea, 50 wt% sucrose, 10 wt% Triethanolamine, and dH2O). Vials containing brains in 1/2-PBS-diluted Reagent-2 were placed in a vacuum desiccator with gentle shaking overnight at RT. The following day, the solution was replaced with Reagent-2 and incubated at 37°C overnight on a rotator. The next day, Reagent-2 solution was replaced with fresh Reagent-2 and incubated at 37°C overnight on a rotator. This step was repeated 4 times. We imaged and processed a total of n = 15 from 5 female and 10 male mice.

### Lightsheet imaging

Samples were imaged within 7 days of completing the clearing protocol at the Gladstone Institutes Histology and Light Microscopy Core on a Zeiss Lightsheet Z.1 microscope. Lightsheet imaging of a mouse P0 brain required approximately 50 tiles (5x10) per brain at 5x (Zeiss EC Plan-Neofluar 5x, NA 0.16, RI 1.45, WD 5.6mm), which afforded a level of resolution that allowed for nuclei detection (0.913 $\mu m$/px lateral resolution).

The microscope requires specimen to be suspended between fixed illuminator and detector objectives, typically using capillaries to embed small specimen. To suspend the relatively larger whole brain, we designed a custom 3D-printed rod with a platform to maximize surface area for gluing the ventral surface of the brain to the mount (Suppl. Fig. S9). The rod also contains a neck to position the brain directly under the mount holder, allowing full movement of the brain throughout the chamber to capture the brain it its entirety. The mount oriented the brain axially toward the detector to minimize the path of the illuminators on opposite sides laterally through the tissue as well as the emission path superiorly to the detector. We designed the mount in Onshape and printed it using a Stratasys uPrint 3D printer.

To glue the brain to the mount, we placed the brain, ventral surface facing up, on a custom sieve to drain reagent and rolled cotton bud applicators on the ventral surface of the brain to dry it. We applied glue (Scotch Super Glue Liquid) with a brush applicator to the mount platform surface and attached it the brain. After flipping the brain to dorsal side up, we placed the mount in a custom holder to allow the glue to dry over 3 minutes and dribbled Reagent-2 media on the dorsal surface to ensure that it did not dry out. After the glue dried, we suspended the mount from the microscope manipulator before immersing the mounted brain into the microscope chamber filled with Reagent-2.

With the lasers initiated, we aligned the two opposite illuminators along the z-axis by visual inspection at a central region of the brain. We ranged the z-stack from just beyond the first and last z-planes with visible nuclei, typically 800-1000 planes using a slice interval of 4.935 $\mu m$ with a lightsheet thickness of 10.44 $\mu m$, and set the image tiling to include the farthest lateral and anterior-posterior nuclei with a 10% overlap per tile. We illuminated the brain with 488nm excitation and 30ms dwell time through the Z.1 LSFM 5x/0.1 illuminators, LBF 405/488/561/640 laser blocking filter, SBS LP 560 secondary beam splitter, and BP 505-545 band pass filter. The microscope paused for 20s between each tile to allow tissue settling after repositioning for the next tile. We controlled the microscope through the Zeiss Zen microscopy software suite and saved images in a CZI multi-tile format with all tiles stored in a single archive.

### Image stitching

To stitch the tiled microscopy images in an automated fashion, we used the Fiji/ImageJ [83], [84] BigStitcher plugin [85], a successor to the Stitching plugin [101] that allows for better memory management and multi-processing as well as a graphical interface to verify alignments. As we needed to stitch multiple brains, we accessed this plugin headlessly through its scriptable interface, manually intervening only to visually verify alignments before proceeding with the fusion step.

After importing the CZI file into the BigStitcher HDF5-based format, the plugin auto-detected the brightest illuminator for each tile, discarding planes from the other illuminator. We chose to simply select planes from the optimal illuminator rather than fusing planes from both illuminators after our inspection revealed that illuminators rarely if ever aligned perfectly throughout the tile, leading to artifacts such as apparent elongation of nuclei from imperfect overlap of the same nuclei from different illuminators.

The plugin calculated tile shifts using a phase correlation method, and we filtered out links below a correlation threshold of r = 0.8 before applying shifts through the two-round iterative global optimization strategy. To account for occasional tile misalignments, we manually inspected every pre-stitched brain to reposition any misaligned tiles, which occurred in  10% of brains. Once tiles aligned, the plugin fused them into a single large TIFF image per channel. We imported this fused file via Python-Bioformats and Javabridge, libraries that allow access to life science formats via Bio-Formats, to a Numpy array format for image processing in our Python-based software as outlined below.

### Automated nuclei detection

We detected nuclei in cleared mouse brains using a 3D Laplacian of Gaussian blob detection technique throughout each whole brain. To perform detections in large images several hundred gigabytes (GB) to over a terabyte (TB) in size, we subdivided the image into many smaller chunks to reduce RAM requirements and maximize parallel processing. We loaded images through the Numpy library's memory mapped method (`load` with the `mmap_mode` option) to load only the necessary parts of the image on-the-fly, allowing us to load small images a chunk at a time without reading the entire volumetric image into memory. We divided the image shape into overlapping chunks to ensure that nuclei at borders would not be missed, with overlap size of approximately the nucleus diameter. After determining the offset and shape of each chunk, we set the image array as a class attribute, initiated multiprocessing (the `multiprocessing.Pool` in the standard Python library), and accessed each chunk as a view in a separate process via class methods to avoid duplicating arrays in memory. Thus, we could control total memory usage by the size of chunks and the number of CPU (central processing unit) cores available for separate processes.

3D cell detection poses a number of challenges including adapting to local variation such as staining inhomogeneity, background variation, and autofluorescence, in addition to overlapping cells in dense tissue [102]. To address these issues, we analyzed images in a local manner by further subdividing each chunk for preprocessing based on its immediate surroundings. We split each chunk into sub-chunks using the same approach as above but ran each sub-chunk serially within each CPU process. In each sub-chunk, we first clipped intensity values at the 5th and 98.5th percentiles (`percentile` in Numpy) to remove extreme outliers, rescaled the intensities from 0-1, and further saturated signal by clipping the rescaled intensities at the 50th percentile. We next enhanced edges by using unsharp masking with a Gaussian sigma of 8 (`filters.gaussian` in scikit-image) to identify sharp details as the difference between an image and its blurred version (which we amplified by a factor of 0.3) and adding back those details to the original image. We mildly eroded the resulting signal with an erosion filter using an octahedron structuring element of size 1 (`morphology.erosion` with `morphology.octahedron` in scikit-image) to separate out blobs.

To detect blobs, we implemented the 3D Laplacian of Gaussian blob detector from the scikit-image library (`feature.blob_log`) as a multi-scale interest point operator [103]. We set the minimum and maximum sigma based on the microscopy resolution, with 10 intermediate values, detection threshold of 0.1, and overlap fraction threshold of 0.55 below which duplicated blobs are eliminated. Initially we missed many nuclei positioned above one another in the z-direction, likely because the anisotropy necessitated by the relatively thick lightsheet at 5x in our setup limited resolution in the z-direction. To improve detection along the z-axis, we interpolated the images in this direction to near isotropy before detection (`transform.resize` in scikit-image). The blob detector had a tendency to cluster detections in the bottom and topmost z-planes in each ROI from nuclei visible within the ROI but whose centroids are outside. To avoid this clustering and minimize duplication with adjacent ROIs, we cropped nuclei from these planes on the assumption that they would be captured better in the adjacent, overlapping ROIs.

Overlapping chunks minimized missing nuclei at edges but also necessitated pruning blobs duplicately detected in adjacent chunks. Pruning involves checking for duplicates within all potentially overlapping regions. Since the overlapping portions of the regularly spaced chunks collectively form a grid pattern throughout the full volumetric image, we could limit our search to these grid planes along each axis. After completing detection on the whole image, we first pooled all detected blobs into a single array. Along a given axis of the full image, we determined the boundaries for each overlapping region and all of its blobs. Within each overlapping region, we found all blobs close to another blob by taking the absolute value of the difference between all blobs with one another and finding blobs within a given tolerance in all dimensions. The tolerance was titrated so that the ratio of final blobs in overlapping regions to the next adjacent regions of same volume was about 1:1. For each close pair of blobs found, we replaced both blobs with a new blob that took the mean of their coordinates. To minimize memory usage, we checked smaller groups of blobs against one another until completing all comparisons. We checked overlapping regions simultaneously in multiprocessing along a given axis for efficiency, re-pooled all blobs, and pruned along the next axis to account for blobs that may have been duplicately detected in overlapping chunks along multiple axes, at grid intersections.

Parameters for preprocessing, detections, and pruning steps were optimized through a Grid Search approach, a type of hyperparameter tuning, to check combinations of parameters systematically. To evaluate the accuracy of each set of parameters, two students in our lab generated truth sets of nuclei locations and radii using our serial 2D nuclei annotation tool, taking ROIs of size 42 x 42 x 32 pixels (x, y, z) from representative ROIs of all major brain structures (n = 15 forebrain, 8 midbrain, and 7 hindbrain; n = 2766 nuclei). They separately generated additional truth sets at a slightly lower magnification (4x), size 60 x 60 x 14 pixels (n = 40 ROIs, 1116 nuclei) to increase representation. After detecting nuclei on these images with a given set of parameters, matches between detections and ground truth was determined using the Hungarian algorithm, a combinatorial optimization method to determine optimal assignments between two sets [104], as implemented in `optimize.linear_sum_assignment` in Scipy. After scaling nuclei coordinates for isotropy, we found the Euclidean distances between detected and truth nuclei points through `distance.cdist`, which serves as the cost matrix input to `optimize.linear_sum_assignment` to find optimal pairings between points based on closest distance. We took correctly identified detections, or true positives (TP), as pairings within a given tolerance distance. Unpaired detections or those in pairs exceeding this threshold were considered false positives (FP), and the same for ground truth were false negatives (FN). Since a match for a given nucleus within the ROI may lie outside of it and thus go unseen, we first searched for pairings only within an inner sub-ROI, followed by a secondary search for pairings between only unmatched inner sub-ROI objects and the rest of the ROI [105]. This approach reduced the total number of nuclei available (n = 1118 nuclei) but avoided missed border matches. As measures of performance of our detection compared with ground truth, we used the following standard equations:

$$\text{Sensitivity (Recall)} = \frac{TP}{TP + FN} \tag{6}$$

$$\text{Positive Predictive Value (PPV, or Precision)} = \frac{TP}{TP + FP} \tag{7}$$

### Image downsampling and registration

To assign nuclei to the proper brain label, we employed automated label propagation by registering the E18.5 atlas to each of our imaged mouse brains using SimpleElastix [68], a toolkit that combines the programmatic access of SimpleITK [76] to the Insight Segmentation and Registration Toolkit (ITK) [75] with the Elastix [65], [66] image registration framework. Elastix has been recently validated as a computationally efficient and accurate tool for registration of mouse brains cleared by CUBIC [67].

As stitched images are typically several hundreds of GBs per file, image downsampling was necessary to reduce memory utilization during image registration. To reduce file size efficiently in both time and memory usage, we employed the same chunking strategy as used during nuclei detection except with larger, non-overlapping units to resize multiple sections of the image simultaneously. We also reduced memory required for the output array by saving directly to disk with a memory-mapped array (`lib.format.open_memmap` in Numpy). We matched the target final size to the E18.5 atlas, which would be registered to each down-sampled image.

Registration involved rigid followed by non-rigid alignment. For rigid registration, we employed a translation (`translation` parameter map in SimpleElastix) with default settings except increasing to 2048 iterations (`MaximumNumberOfIterations` setting) followed by an affine (`affine` parameter map) with 1024 iterations, applied with an `ElastixImageFilter`, to shift, resize, and shear the atlas microscopy image to the same space as that of the sample brain. For non-rigid alignment, we employed a b-spline strategy (`bspline` parameter map) guided by the `AdvancedNormalizedCorrelation` as the similarity metric [67], [69] with grid spacing of size 60, measured in voxels rather than physical units (`FinalGridSpacingInVoxels` setting in place of `FinalGridSpacingInPhysicalUnits`), over 512 iterations. The `TransformixImageFilter` in SimpleElastix allowed us to apply the identical registration transformation to the atlas labels image, except that we set the final b-spline interpolation order (`FinalBSplineInterpolationOrder`) to 0 to avoid interpolating any new values, preserving the labels' specific set of integer values. We applied this identical transformation to both the mirrored and edge-refined atlas labels.

To evaluate the level of alignment from registration, we measured the similarity between each registered atlas histology and its corresponding sample image using a Dice Similarity Coefficient (DSC) [51] as implemented by the `GetDiceCoefficient` function in SimpleElastix/SimpleITK, given by the equation [52]:

$$\text{Dice Similarity Coefficient (DSC)} = 2\frac{|S \cap T|}{|S| + |T|} \tag{8}$$

where S and T are two different sets of voxels. We took the foreground of each atlas and sample microscopy image to be its mean threshold (`filters.threshold_mean` function in scikit-image) and input them to a `LabelOverlapMeasuresImageFilter` to take the DSC.

### Whole brain nuclei measurements by label

Registration of the atlas to our volumetric nuclear-stained brain microscopy images allowed us to quantify nuclei per label for comparison with the original (mirrored) and smoothed (edge-aware) atlases. We first measured volumes per label by taking a mask of each registered label and summing the foreground pixels within each mask before multiplying this volume by the microscopy pixel resolution (scaled for downsampling) to obtain volumes in physical units ($\mu m^3$).

For nuclei densities, we first constructed a nuclei heat map by converting the nuclei coordinates to nuclei per voxel within the downsampled image. We scaled the coordinates to the scaling of the downsampled image, rounding to the nearest integer, and found the counts of nuclei at each coordinate (`unique` with `return_counts` option in Numpy). We next indexed these coordinates directly into an empty Numpy array of the same shape as that of the downsampled image to assign them to the corresponding nuclei counts at each voxel. We used the same label mask previously obtained to find the number of nuclei within each given label. Dividing the number of nuclei by the volume within each label gave the label nuclei density.

To measure the variability of nuclei within each label before and after label reannotation, we measured the coefficient of variation within each label given by the standard equation:

$$\text{Coefficient of variation (CV)} = \frac{\sigma}{\mu} \quad (9)$$

where $\sigma$ is the standard deviation, and $\mu$ is the mean. A lower coefficient of variation indicates lower variability and thus tighter capture of a more homogeneous label. As a raw proxy for nuclei variation, we first measured the variation of intensities within the nuclear-stained images. Using the same label masks, we took the standard deviation of voxel intensities and divided it by the mean of intensities within the label (`std` and `mean`, respectively, in Numpy) to obtain the intensity coefficient of variation. Similarly, we measured the nuclei coefficient of variation by measuring the standard deviation and mean values of nuclei counts per label within the nuclei heat map.

### Nuclei clustering

As another measure of label alignment at the nuclei level, we measured nuclei clustering using Density-Based Spatial Clustering of Applications with Noise (DBSCAN) [70] as implemented by the scikit-learn library [82]. DBSCAN clusters tightly packed points within a neighbor distance defined by the parameter $\epsilon$ and a minimum number of points given as another parameter, with isolated points in lower density regions that cannot be clustered considered "outliers" or "noise." The minimum number of samples is typically taken as $2 \cdot ndim$, where $ndim$ is the number of dimensions [106], thus giving 6 for our 3D nuclei point cloud. To find $\epsilon$, the nearest-neighbor distance of the $2 \cdot ndim - 1$ neighbor for each point is sorted and plotted to find the distance at the "elbow" point, the point of maximum curvature [106], which we found to be at least $20\mu m$ (Suppl. Fig. S14A, B).

For each label in the original (mirrored) atlas registered to each wild-type brain, we extracted the nuclei coordinates within the label and clustered them by DBSCAN to find the number of clusters, nuclei per cluster, and "noise," or number of isolated nuclei that remained unclustered. We repeated the same process using the same nuclei coordinates for each brain but with the smoothed (edge-aware) atlas registered identically to the brain. As the "elbow" distance of maximum curvature can be difficult to define, and $\epsilon$ can strongly influence the clustering, we repeated this process for a range of $\epsilon$ values through the elbow region (Suppl. Fig. S14C) and highlighted for a conservative distance of 20 (Suppl. Fig. S14D, E).

### MagellanMapper software suite

We provide the MagellanMapper image software suite as a tool to assist with visualization, annotation, and automated processing of volumetric images (Suppl. Fig. S15). The suite consists of a graphical user interface (GUI) to aid visualization of 2D images in a 3D context and command-line interface (CLI) for non-interactive processing in workstation and cloud environments.

#### Graphical interface

The main GUI integrates ROI selection with 3D point and surface rendering through the Mayavi toolkit [78]. Users can load volumetric image files and specify ROI boundaries through sliders and text boxes or load a previously saved ROI. 3D point rendering provides a voxel-based visualization of the ROI with minimal filtering, whereas the 3D surface rendering utilizes VTK (The Visualization Toolkit) [107] for cleaner images. The interface is implemented in TraitsUI for integration with Mayavi.

To inspect raw images, the user can launch two types of mixed 2D/3D interfaces from the main GUI to display and annotate the original 2D images for each plane. The first 2D interface is a serial 2D ROI viewer that shows each successive 2D plane within the ROI side-by-side, allowing the user to follow objects such as nuclei that come and go from plane to plane. Larger overview images at different magnifications show context and synchronize with the smaller views to effectively zoom in on a given plane. These overview images are also scrollable along the z-axis to visualize subtle object shifts in-place. The interface also provides annotation tools geared toward blob detection such as nuclei. Detected blobs appear as circles along with optional segmentations, and the user can drag, resize, or cut/copy/paste circles to improve placement and flag their correctness. We have used this simplified blob annotator to generate truth sets for blob detection verification and optimization.

In addition to ROI viewers, we provide a simultaneous orthogonal viewer to visualize and annotate atlases in all three dimensions. It displays orthogonal planes of the full volumetric image in three separate panels, with crosshairs in each panel denoting the corresponding planes in other panels. Clicking on or scrolling within any panel updates crosshairs and synchronizes the other displayed planes. The user can also load label maps to overlay directly on atlas microscopy images with an adjustable labels opacity to allow close inspection of annotation alignment with anatomical structures. To distinguish an arbitrary number of labels from one another, we use a custom discrete colormap with randomly generated colors, each assigned to a single label. We display all images as views of one another to minimize memory requirement and update all orthogonal planes in real-time.

As a method for simple, rapid editing of these labels, the user can enter an editing mode to simply click and drag on individual labels to paint them into other spaces. We have used the interface with a tablet and electronic pen to edit labels by drawing. As hand-editing any given plane likely introduces edge artifacts seen in other orthogonal directions, we also designed a method to interpolate contours between two distant planes. After the user edits the same label ID at start and ending planes and initiates the interpolation, it takes the signed distance transforms of the label masks (`ndimage.distance_transform_edt` in Scipy combined with a mask of the original label to identify distances inside versus outside the label) and interpolates those distances for each intervening plane (`interpolate.interpn` applied across a meshgrid of the planes). The interpolation provides label border extensions that are generally smooth in all dimensions after manually editing only the first and last plane.

The annotation interfaces are implemented in Matplotlib [77]. Blobs are stored in an SQLite [108] database, while atlases edits are saved directly to their underlying 3D image file.

#### Headless pipelines

In addition to a GUI for interactive visualization and verification, MagellanMapper provides automated pipelines for non-interactive image processing such as whole brain nuclei detection in cloud-based work environments. Users can access the suite via its CLI, and the suite provides Bash scripts to connect MagellanMapper with other tools such as Fiji/ImageJ for image stitching and Amazon Web Services (AWS) in a platform-independent manner.

For input/output (I/O), the suite utilizes standard 3D image formats for portability with other software libraries. Microscopy images stored in proprietary formats such as Zeiss CZI format can be imported to a standard Numpy array archive using the Bio-Formats library [109] (via the Javabridge and Python-Bioformats libraries developed for CellProfiler [110] libraries) along with a separate Numpy archive containing image metadata extracted from the original file. Loading the imported Numpy array as a memory-mapped file (`load` function with `mmap_mode` option) allows users to access small parts of large files to minimize load time and memory usage by only loading the requested ROI rather than the potentially TB-sized full volumetric image. In addition to Numpy array archives, other 3D image formats such as MetaImage, NifTI, and DICOM are supported through the SimpleITK/SimpleElastix library. Annotated images are saved to their respective formats.

As an example of an automated pipeline for image import, a user can launch the script from a cloud-based server instance to first retrieve and decompress an archived microscopy file previously saved in a cloud storage location. After extracting the microscopy image, the pipeline script launches Fiji/ImageJ to run a custom headless script for the BigStitcher plugin, which stitches the tiled microscopy image as described above. After allowing the user to verify and adjust tile placements through the BigStitcher interface, the script continues the BigStitcher tile fusion operation, resulting in TIFF formatted images for each channel. The pipeline script next calls MegellanMapper to import these TIFF images into a single Numpy array archive. Subsequent pipelines can be run to process the imported image for automated nuclei detections, downsample or transpose the image, register images to an atlas, or automatically refine new atlases in 3D.

#### Software access

We provide MagellanMapper as open-source software in the hope of facilitating both interactive and headless processing of large volume microscopy images and the atlases to which they will be registered. The suite API includes 3D image processing functions designed to be useful as library methods for other applications. Library functions for the Python and R plots depicted here are also provided to reproduce similar graphs. As an open-source, Python-based tool, our vision for the suite is that it will work alongside, integrate with, and itself become refined by the many other excellent image processing software suites and libraries available to the scientific community.

### Software versions

The MagellanMapper suite is written in Python with the following versions and Python libraries: Python 3.6, Numpy 1.15, Scipy 1.1, scikit-image 0.14, scikit-learn 0.21, Matplotlib 3.0, Matplotlib ScaleBar 0.6, Mayavi 4.6, TraitsUI 6.0, PyQt 5.11, Python-Bioformats 1.1, Javabridge 1.0, SimpleElastix 1.1, and Pandas 0.23. We used Conda 4.7 for Python library management. We stitched images with Fiji/ImageJ 1.52 using BigStitcher 0.2.10. We computed additional statistics in R 3.5 with RStudio 1.1. To interact with AWS, we used AWS CLI 1.16 and Boto3 1.9. For image acquisition, we used Zeiss Zen 2014.

We conducted most software development on a Mid-2014 MacBook Pro (Intel Core i7-4980HQ, 16GB RAM, 1TB SSD) with MacOS 10.13, image stitching and nuclei detections on AWS EC2 c5.9xlarge instances (36 vCPUs, 72GB RAM, typically configured with 1TB SSD) running RHEL 7.5 and Ubuntu 18.04, and volume measurements and aggregation on a Dell Precision T7500 workstation (64GB RAM, 256GB SSD and 2.2TB HDD) with Ubuntu 18.04. We conducted additional cross-platform compatibility testing on a Microsoft Surface Pro 5 laptop (8GB RAM, 256GB SSD) with Microsoft windows 10 (build 1803), Windows Subsystem for Linux running Ubuntu 18.04, and Windows 10 virtual machines in VirtualBox 6.0.

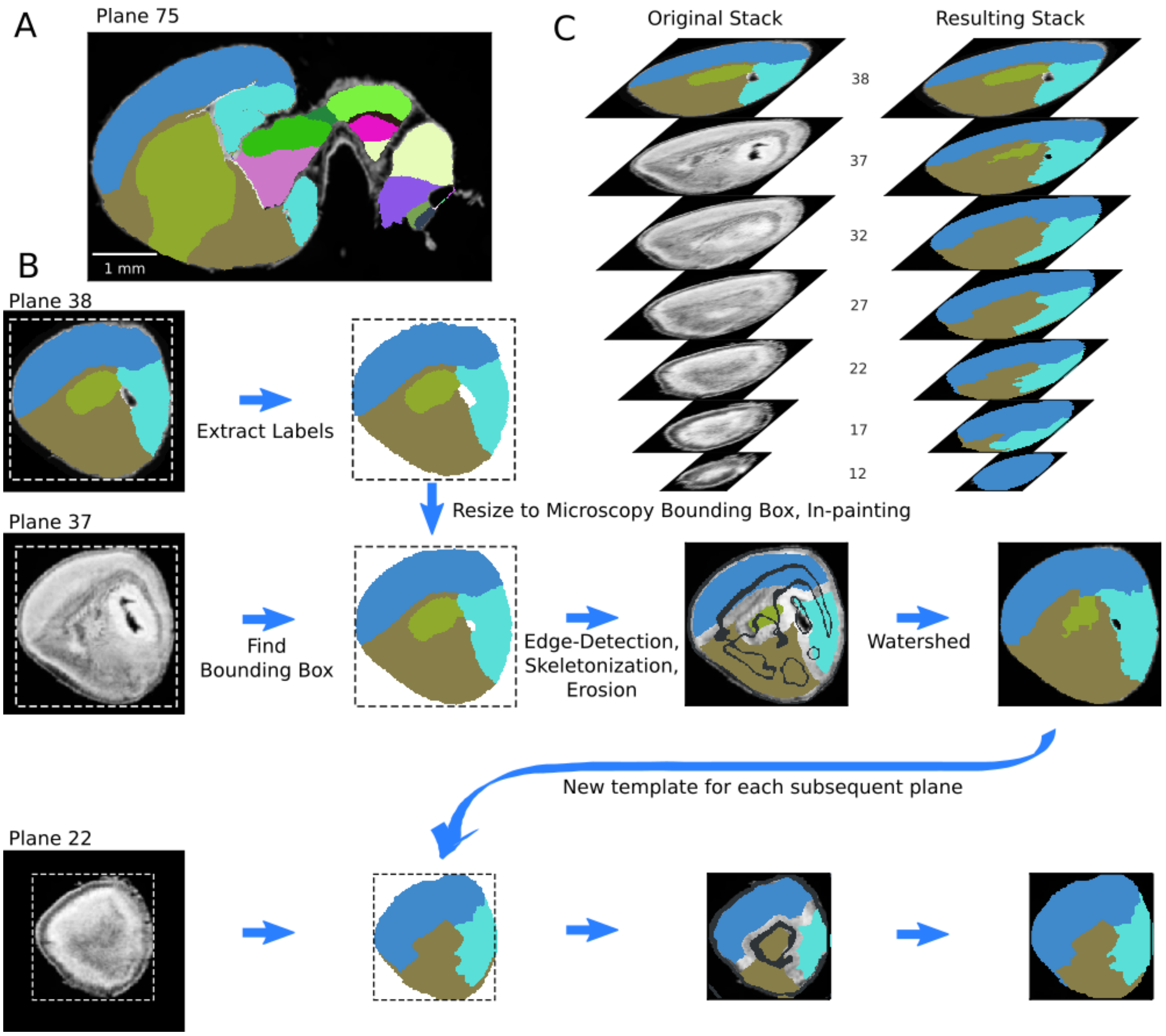


Figure S1: **Lateral edge extension** (A) Example labeled sagittal section toward the lateral edge in the ADMBA E18.5 atlas. (B) Overview of the extension algorithm. Plane 38 represents one of the farthest lateral labeled planes. For each discrete structure, the labels are extracted to serve as templates for the subsequent plane. In plane 37, the corresponding template is resized to the bounding box of the microscopy image in that plane. To further fit labels to the underlying anatomy, gross anatomical edges are found in 3D for the full volumetric microscopy images. Labels are eroded, skeletons are added back to avoid loss of thin sections, and labels are regrown by a compact watershed guided by the anatomical edges in the plane. This refined plane of labels thus becomes the template for the next plane. As the extension progresses, some labels disappear during the erosion step, preferentially central labels, modeling the tapering of labels laterally. (C) Every fifth label in a stack after the original template plane in (left) the original atlas and (right) after label extension.

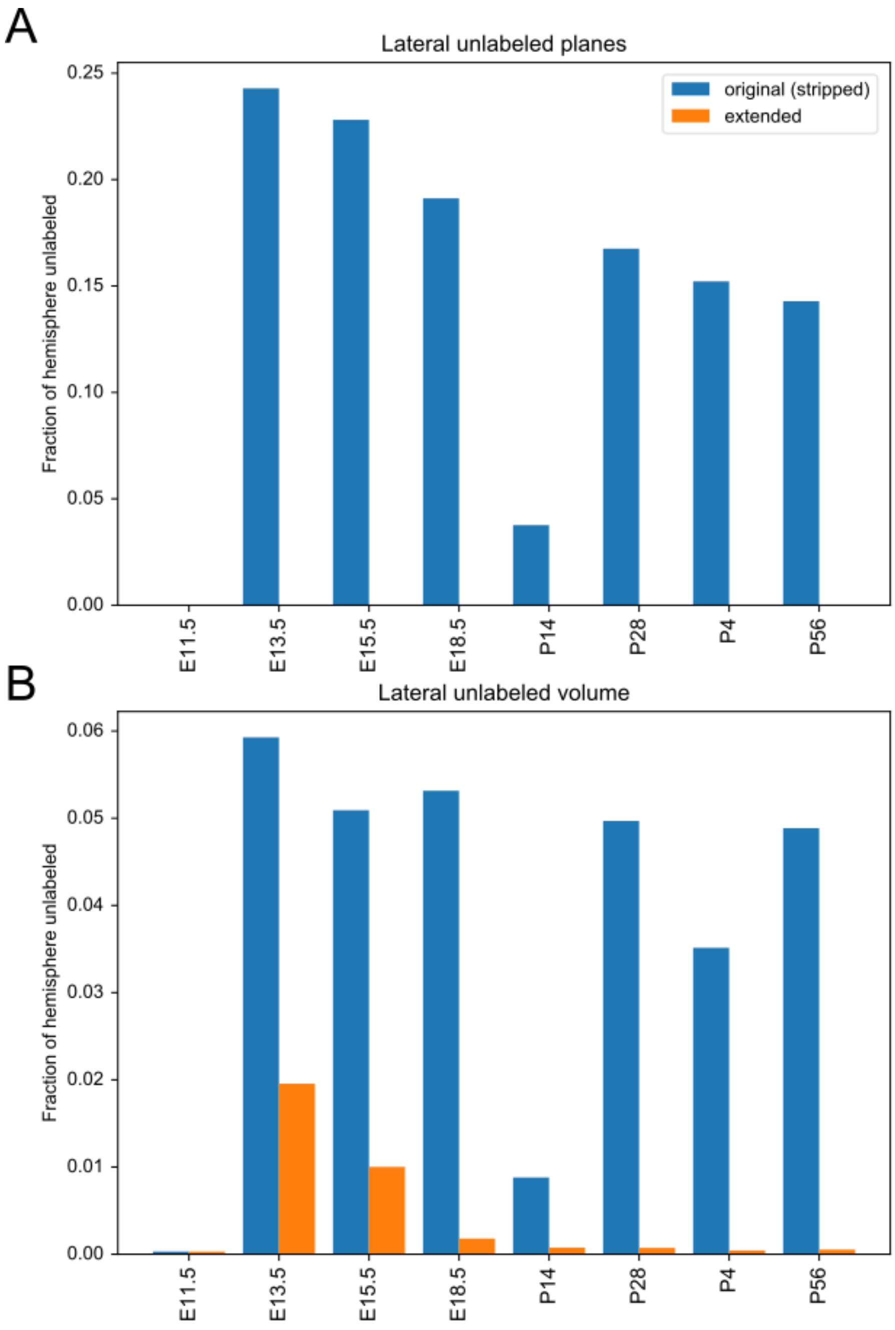


Figure S2: **Fraction of hemisphere that is unlabeled in each of the ADMBA atlases** (A) The fraction of unlabeled planes in a given hemisphere is taken as the number of sagittal planes without any labels over the total number of sagittal planes in the hemisphere that should have labels. By definition the extended atlases have no detected unlabeled planes because any plane determined to require labeling was filled. (B) Unlabeled volume fractions are measured by taking label volume over the thresholded histology foreground in the hemisphere. Each hemisphere is taken as the predominantly labeled hemisphere of the given atlas.

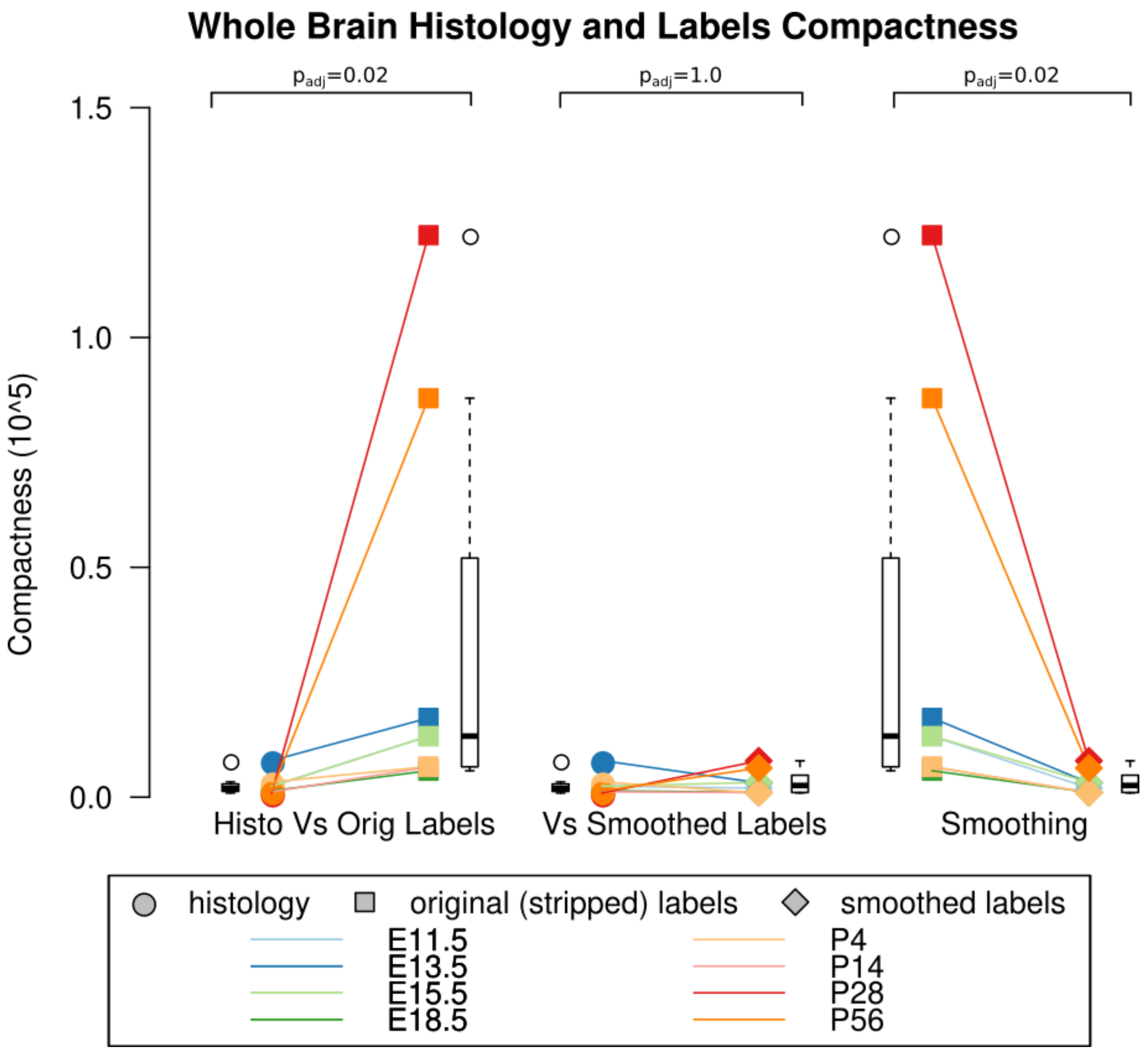


Figure S3: **Compactness of whole brain histology and labels** (A) Compactness of the whole brain histology can serve as a proxy for compactness of a biological structure. This compactness is significantly lower (i.e. more compact) compared with that of the original (stripped) labels ($p = 0.02$, WSRT, Bonferroni corrected across all comparisons) but (B) similar to that of smoothed labels ($p = 1.0$). (C) Original (stripped) were also significantly less compact (higher compactness) than smoothed labels were ($p = 0.02$).

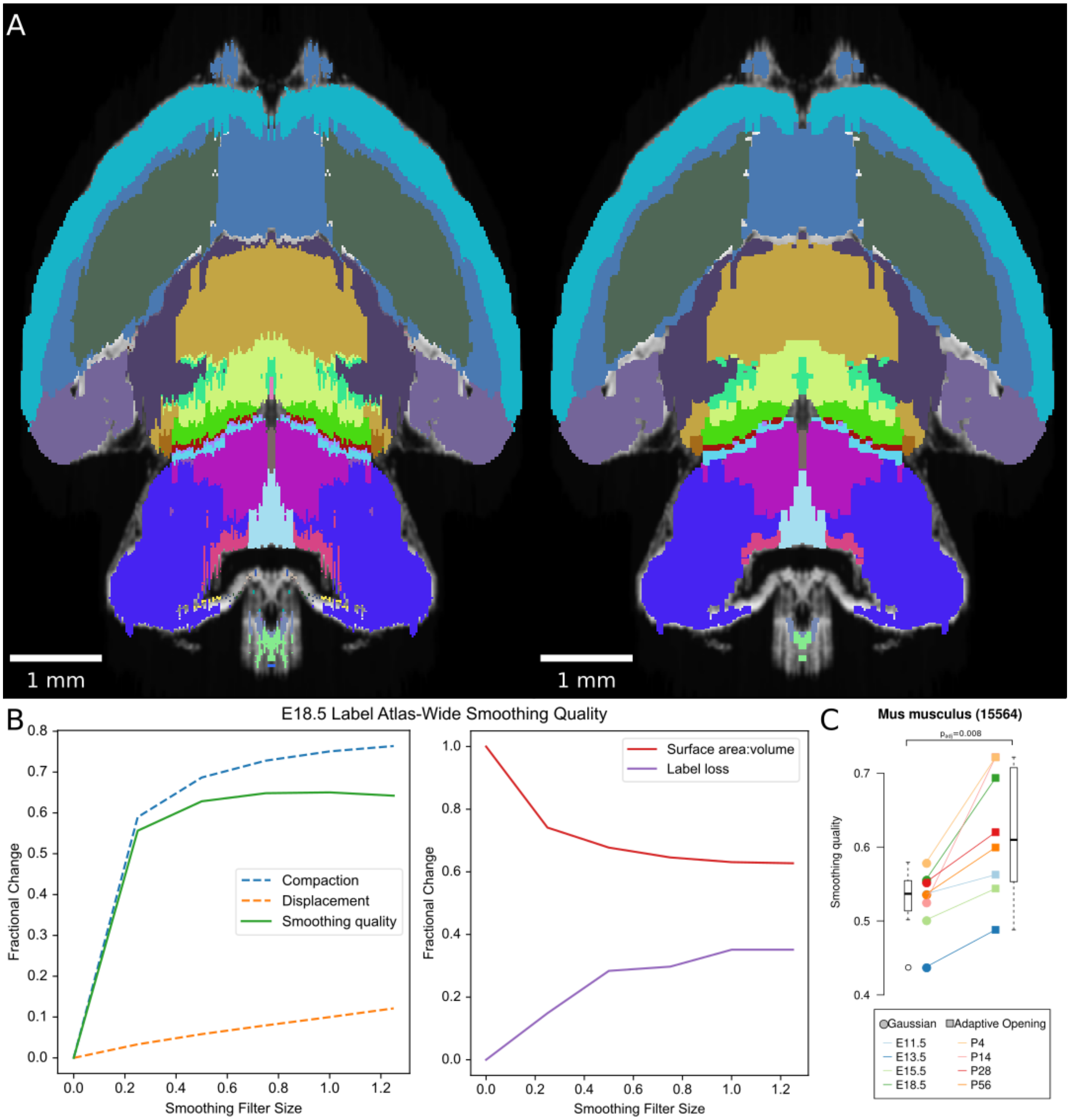


Figure S4: **Smoothing labels by Gaussian blur** (A) Example of the E18.5 atlas before (left) and after (right) smoothing by a Gaussian filter with a sigma of 0.25. (B) Smoothing quality metrics show increased compaction with a relatively slower increase in displacement with increasing filter sizes, leading to a peak smoothing quality at sigma of 1. This smoothing comes at a cost of label loss at all tested sigmas, including 15% at even the lowest tested sigma of 0.25. (C) Compared with Gaussian smoothing at this sigma across all ADMBA atlases, the adaptive opening filter approach showed a significant increase in atlas-wide smoothing quality (median 0.54 by Gaussian vs. 0.61 by adaptive opening filter; p = 0.008, WSRT, Bonferroni corrected; mean 0.53 vs. 0.62).

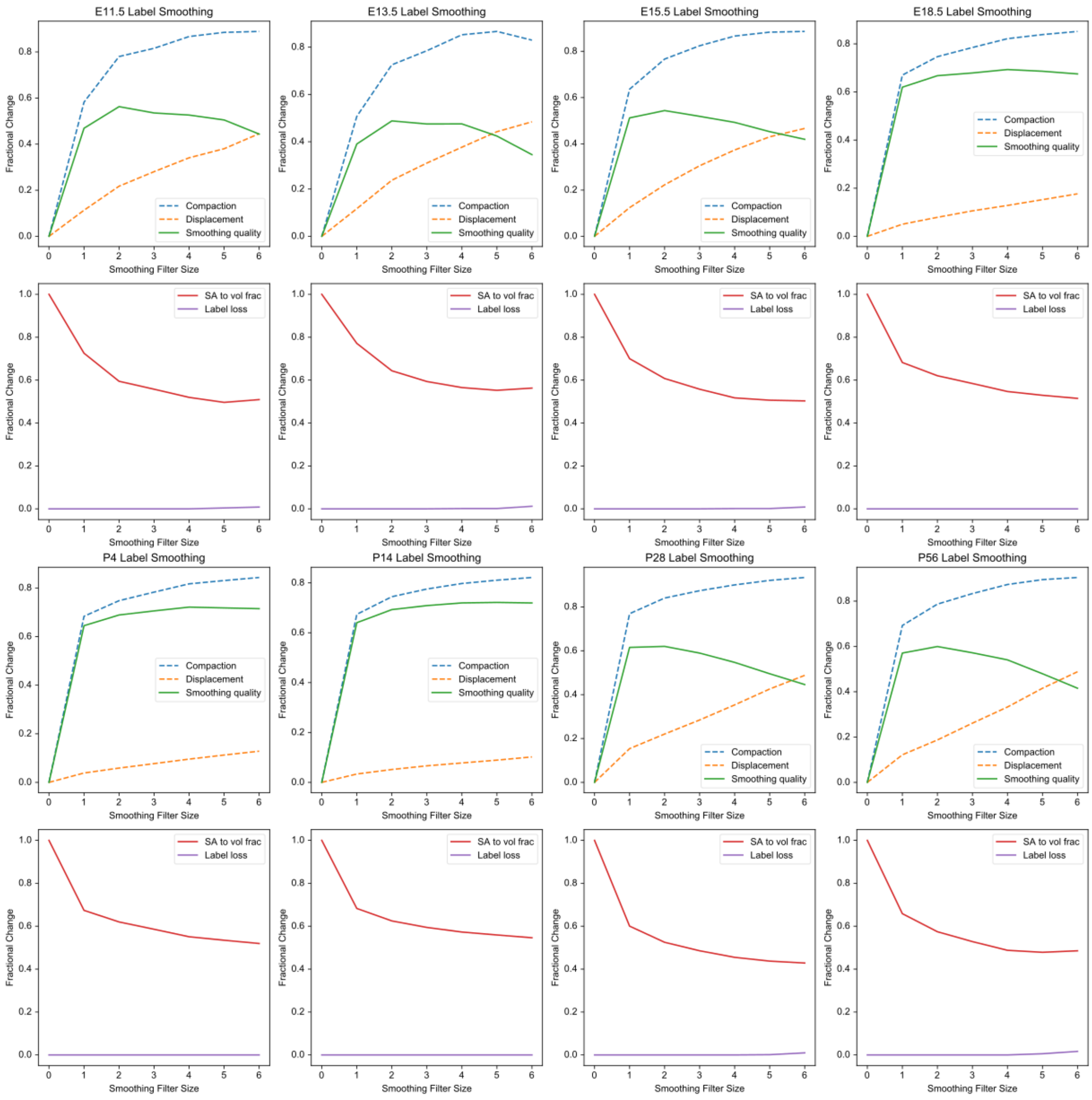


Figure S5: **Smoothing quality across filter sizes for all ADMBA atlases** Smoothing quality metrics for each atlas in the ADMBA across filter structuring elements sizes, with size 0 corresponding to the original (mirrored) atlas. For each atlas, the top plot depicts the overall, brain-wide smoothing quality along with its separate compaction and displacement components, with peak smoothing quality lying between filter sizes 2-6. The bottom plot for each atlas shows the overall surface area to volume across labels as well as fraction of labels lost during smoothing, typically occurring at larger sizes, starting at 4 or higher.

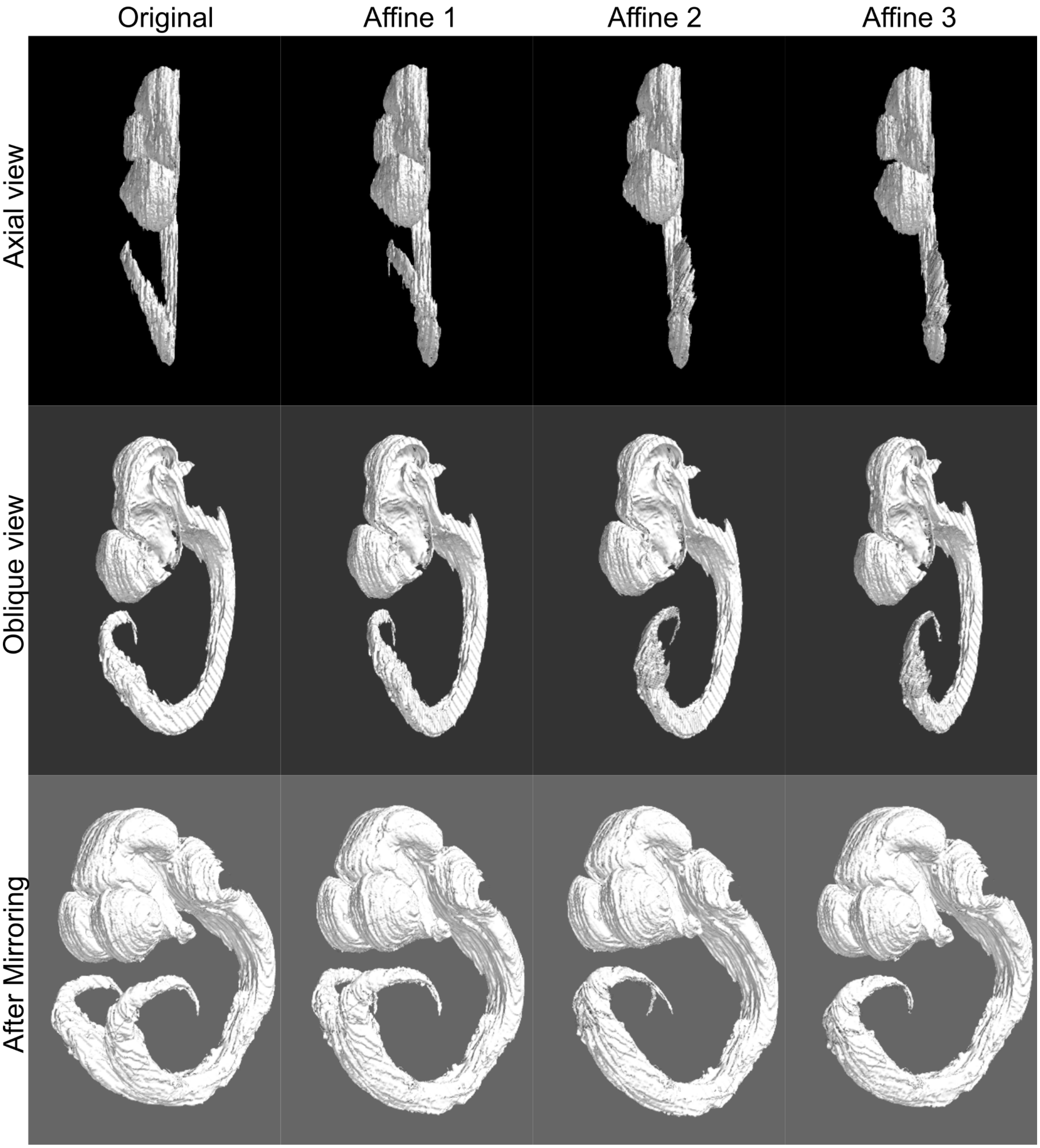


Figure S6: **Piecewise 3D affine transformation of the spinal cord in the E11.5 atlas** (column 1) The distal cord of the ADMBA E11.5 atlas specimen is skewed laterally as seen in the axial view (top), preventing mirroring without duplicating the distal cord (bottom). (columns 2-4) A series of three successive piecewise affine transformations straightened the cord to avoid duplication with mirroring.

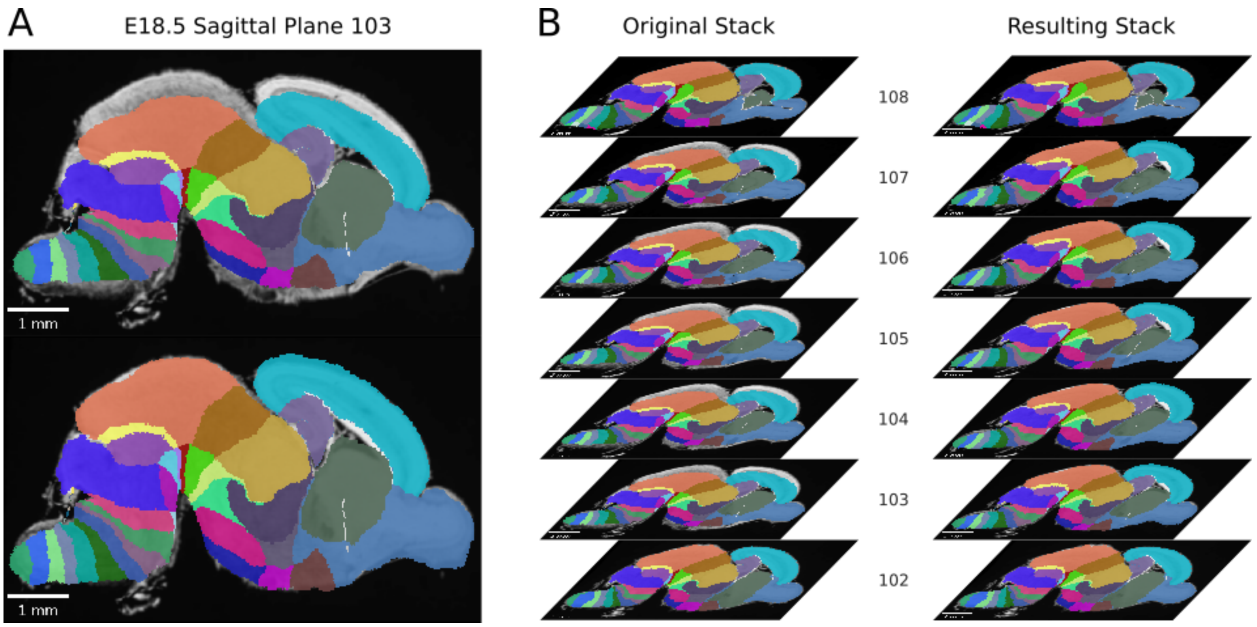


Figure S7: **Expansion of compressed labels in a stretch of planes in the E18.5 atlas** (A) Example plane from a stretch of sagittal planes in the original ADMBA E18.5 atlas where the labels are compressed dorsoventrally. The original labels (top) were re-expanded (bottom) to match the extent of the underlying histology section. (B) The stack of original (left) and re-expanded (right) planes from just before until just after this stretch of compressed label planes.

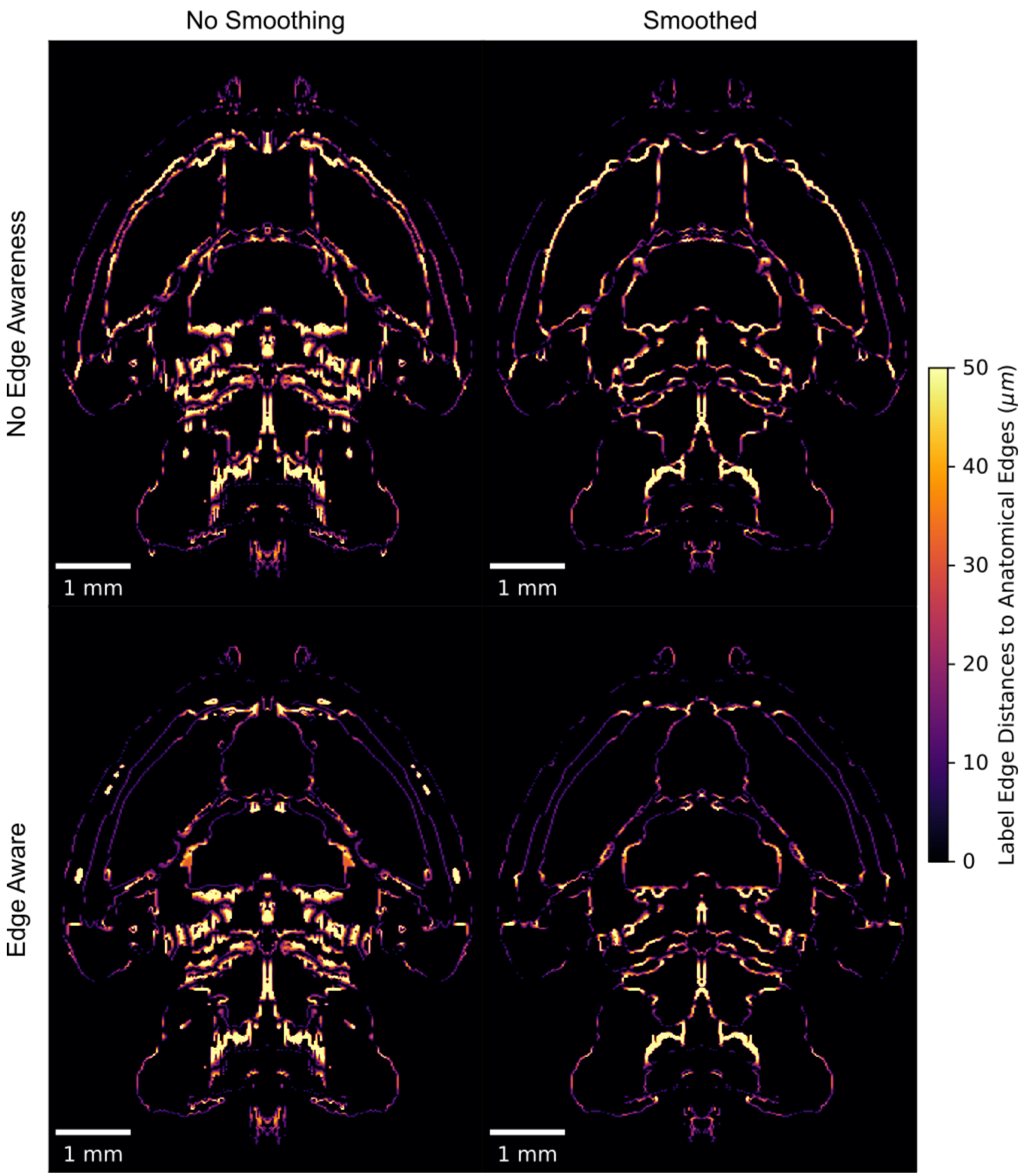


Figure S8: **Edge distances with smoothing, edge-aware reannotation, or both** Distances between labels and gross anatomical edges in the ADMBA E18.5 atlas show improvement from the original (top left) with either smoothing (top right) or edge-aware reannotation (bottom left) alone, but even more so when combining the two approaches through edge aware reannotation followed by label smoothing (bottom right).

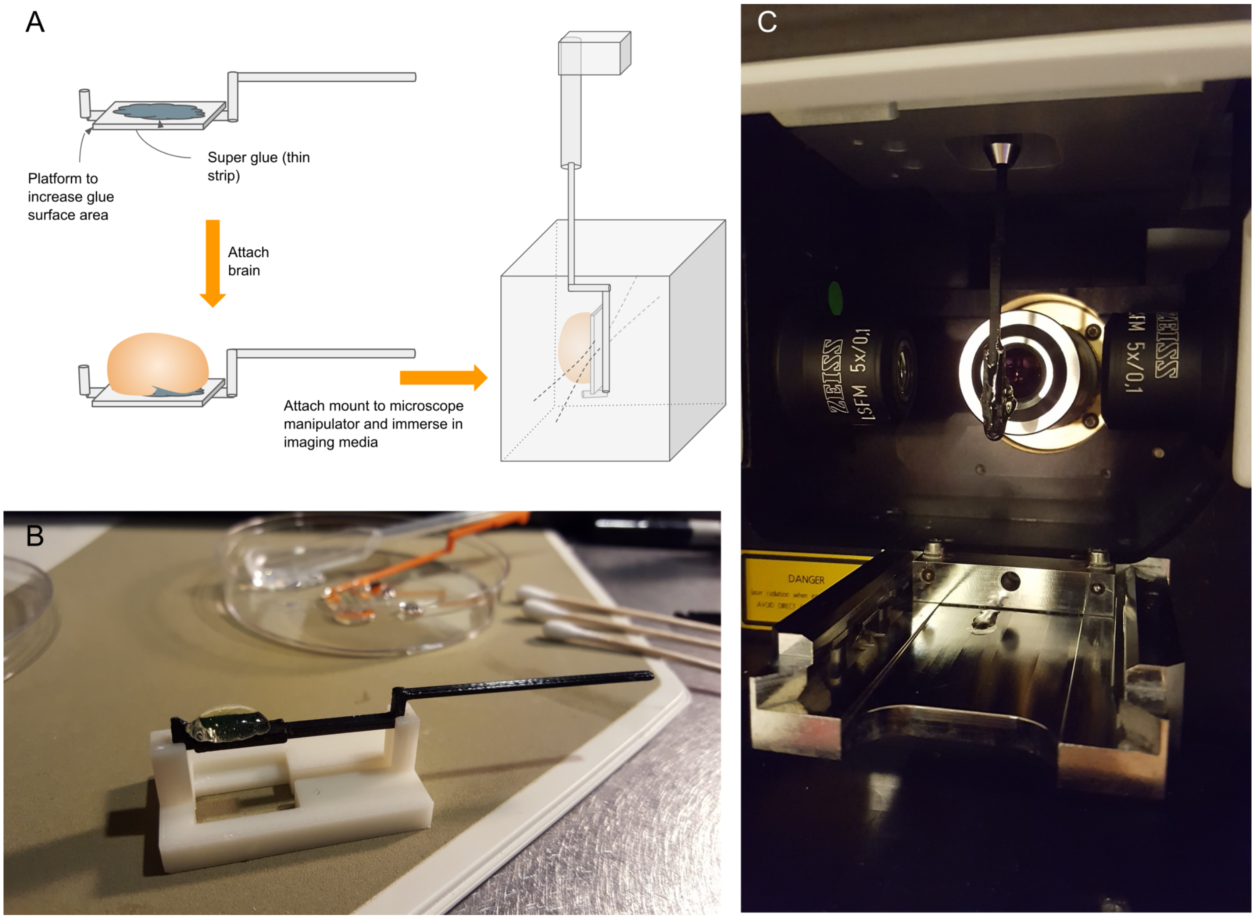


Figure S9: **Custom 3D printed specimen mount for suspending samples in the Zeiss Lightsheeet Z.1** (A) Mount schematic and orientation in the chamber. The lightsheet chamber requires the specimen to be suspended between the two illumination objectives producing a plane of light and detection objective perpendicular to this plane. To hold the brain in place while immersed in the refractive index matching solution, we devised a mount with a platform to maximize surface contact with the ventral surface of the brain, attached by super glue. To allow full latitude of the specimen within the chamber, we added a mount neck to center the specimen between the illumination objectives when the manipulator was approximately centered. (B) Photograph of mount and custom mount holder to facilitate gluing the specimen to the mount platform. Cotton buds were used to dry the ventral surface of the brain before attachment. We used the rod end of another mount to gently press the brain into the glue. (C) Photograph of specimen suspended and centered between the illumination objections, facing the detection objective.

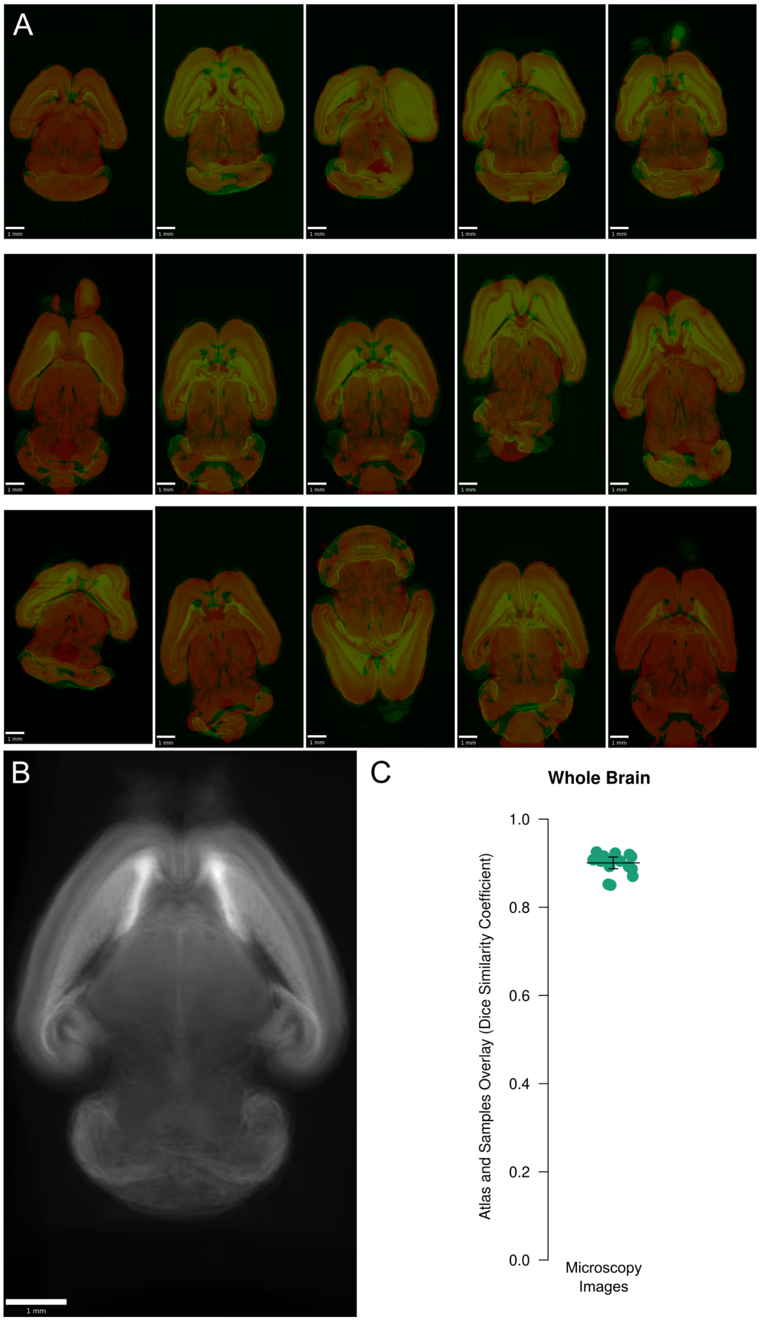


Figure S10: **Registration of atlas to WT brains** (A) Overlays of each registered atlas microscopy image on its corresponding downsampled wild-type brain. All examples are shown at the same plane. (B) Composite image of wild-type sample brains registered in reverse, using the same settings but registering the sample to the atlas to evaluate overall alignment of registered brains. (C) Overall registration quality as expressed by the DSC between the foreground of each sample and its registered atlas showed a median of 0.91 (mean 0.90).

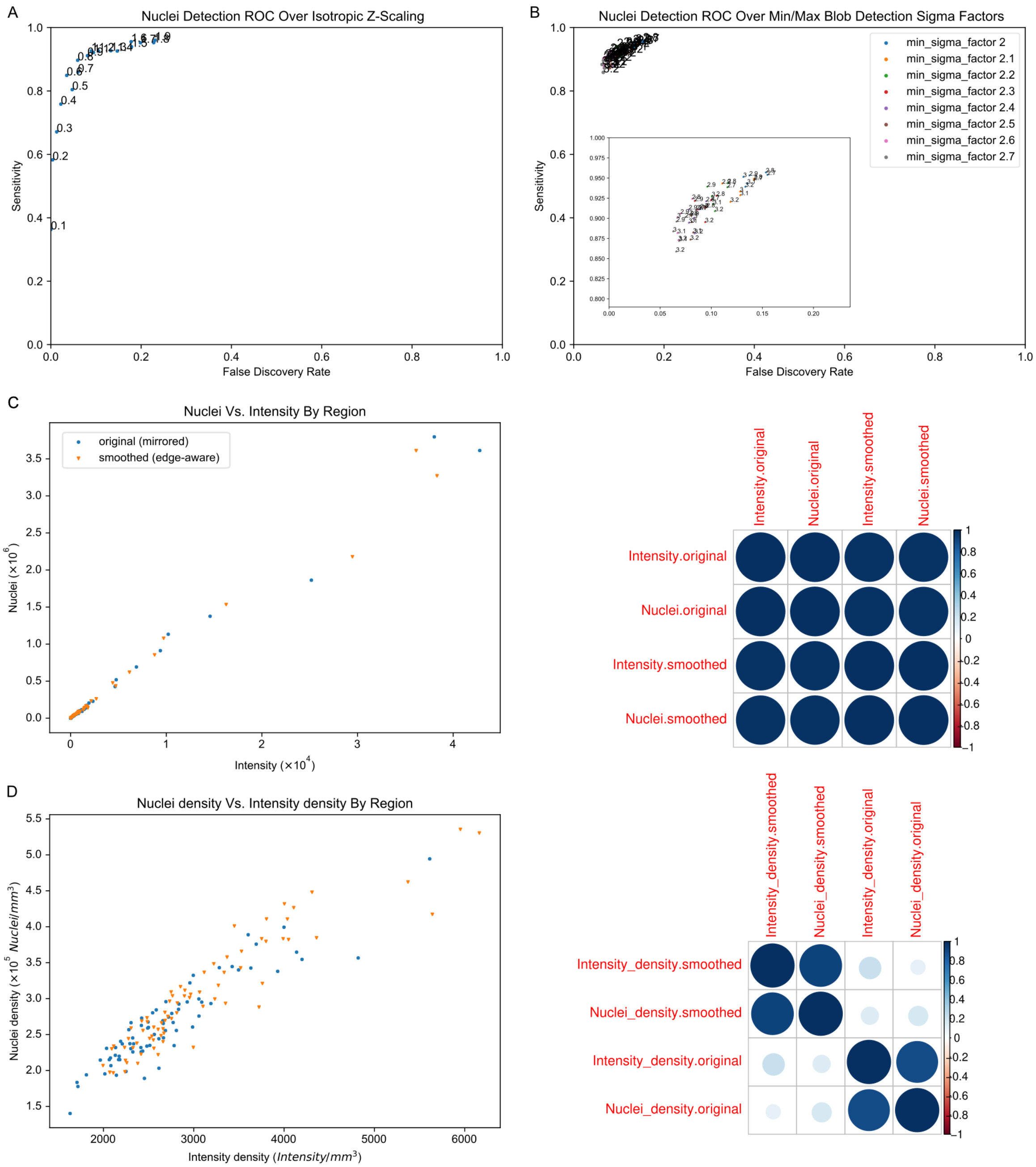


Figure S11: **Nuclei detection optimization and comparison with intensity** (A) Example receiver operating characteristic (ROC) curve of nuclei detection across varying isotropic scaling factors along the z-axis. Scaling is shown as the fraction of isotropy along the z-axis, where 1 = isotropic. For example, 0.5 indicates that each ROI was resized so that the z-axis was half of the size it would need to be for full isotropy with the x- and y-axes. (B) Example ROC evaluating two hyperparameters, the minimum and maximum standard deviations for the Gaussian kernel used for multi-scale detection, to find the optimal size bounds for blob detection. Inset shows zoomed view of the clustered points. (C) Total nuclei vs. intensity by label showed a linear relationship using labels defined by either the original (mirrored; r = 0.997, p $\leq 1x10^{-16}$) or smoothed (edge-aware; r = 0.997, p $\leq 1x10^{-16}$) atlases. In the correlation plot, the size of the circle indicates the size of the correlation coefficient (r). (D) Similarly, nuclei density vs. intensity density relationships were approximately linear in the original (mirrored; r = 0.894, p $\leq 1x10^{-16}$) and smoothed (edge-aware; r = 0.929, p $\leq 1x10^{-16}$) atlases.

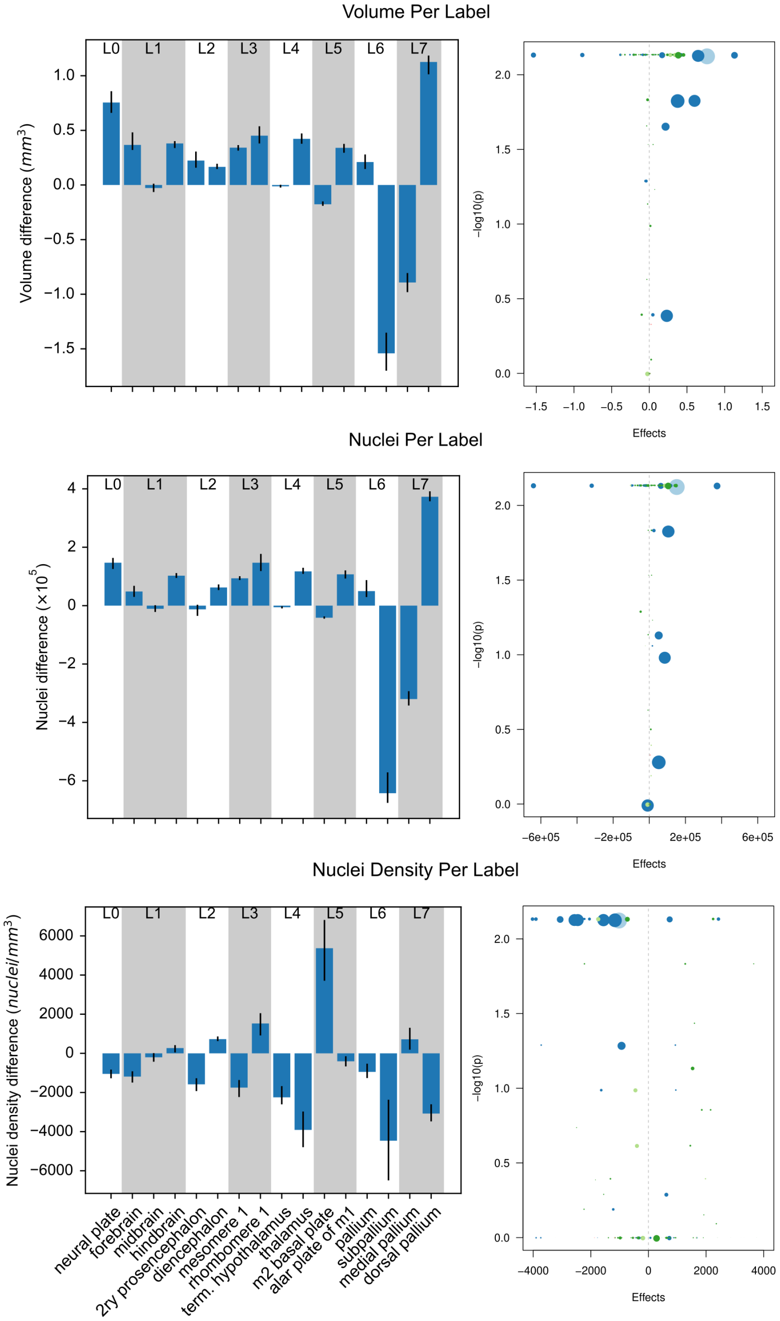


Figure S12: **Changes in volumes, nuclei, and densities across wild-type brains based on atlas labels** The differences between the original (mirrored) and smoothed (edge-aware) labels' volume (top), nuclei (middle), and nuclei densities (bottom) are quantified with bar (right) and volcano (left) plots. Selected regions are shown across the hierarchy of labels from the grossest (neural plate, L0, left) to the finest (e.g. dorsal pallium, L7, right). Positive values indicate decreased size in the refined atlas. Error bars represent 95% confidence intervals. Volcano plots depict each label as a separate point with point size correlating with label size and colors corresponding to major parent structure (neural plate = light blue, forebrain = dark blue, midbrain = light green, hindbrain = dark green). Points higher and farther from 0 have stronger statistical significance and effect size, respectively.

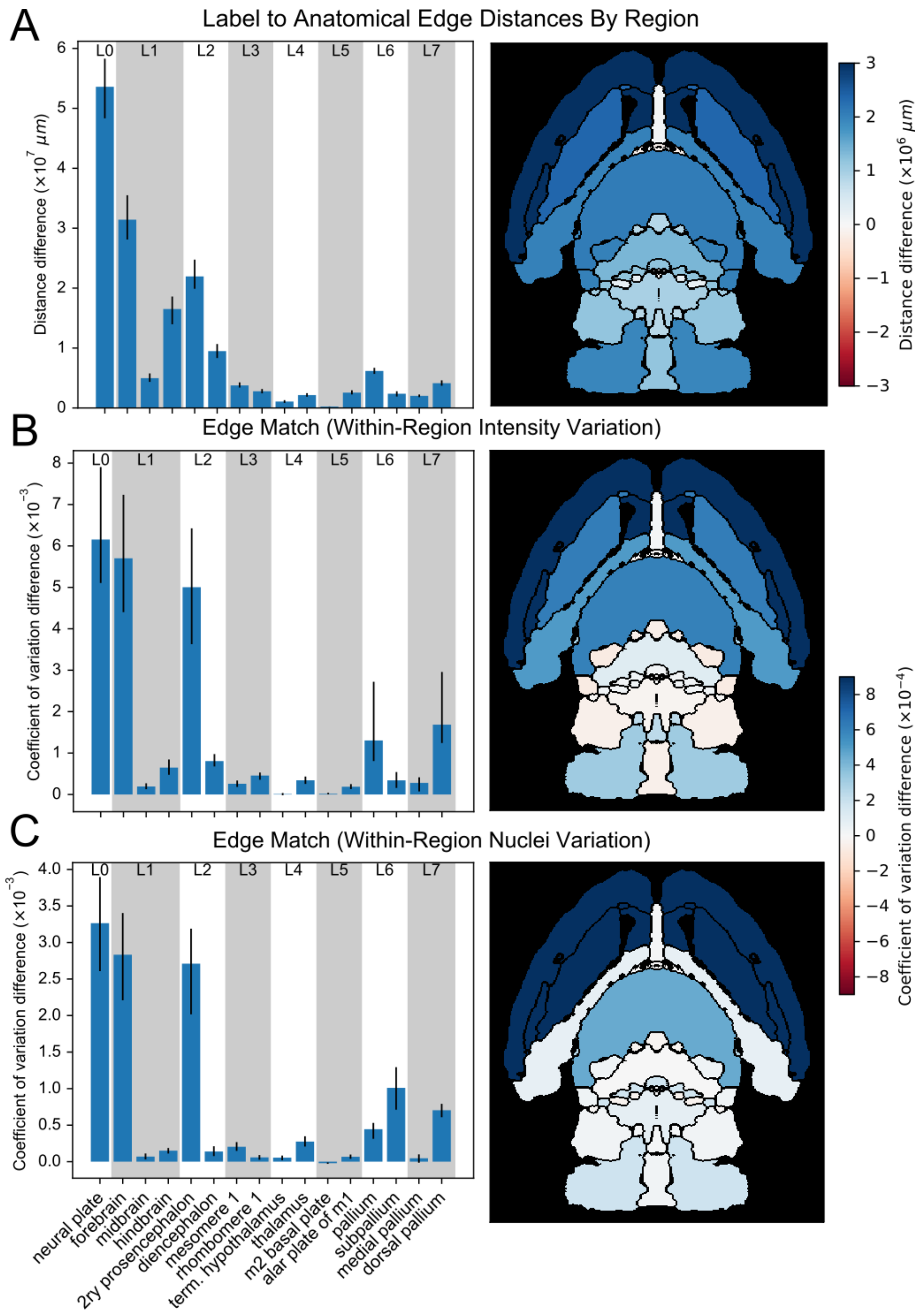


Figure S13: **Region homogeneity in wild-type brains based on the original vs. refined atlases** (left)

The differences between the original (mirrored) and smoothed (edge-aware) labels' edge distance sum (top) and intensity (top) and nuclei (bottom) coefficients of variation as reflections of region homogeneity are quantified with bar plots for selected regions across the hierarchy of labels. Error bars represent 95% confidence intervals. (right) Anatomical maps depict these metrics as a color gradient across all of the finest labels present in a cross section. Increases in a metric with the original atlas are colored in blue, while red represents increases with the refined atlas.

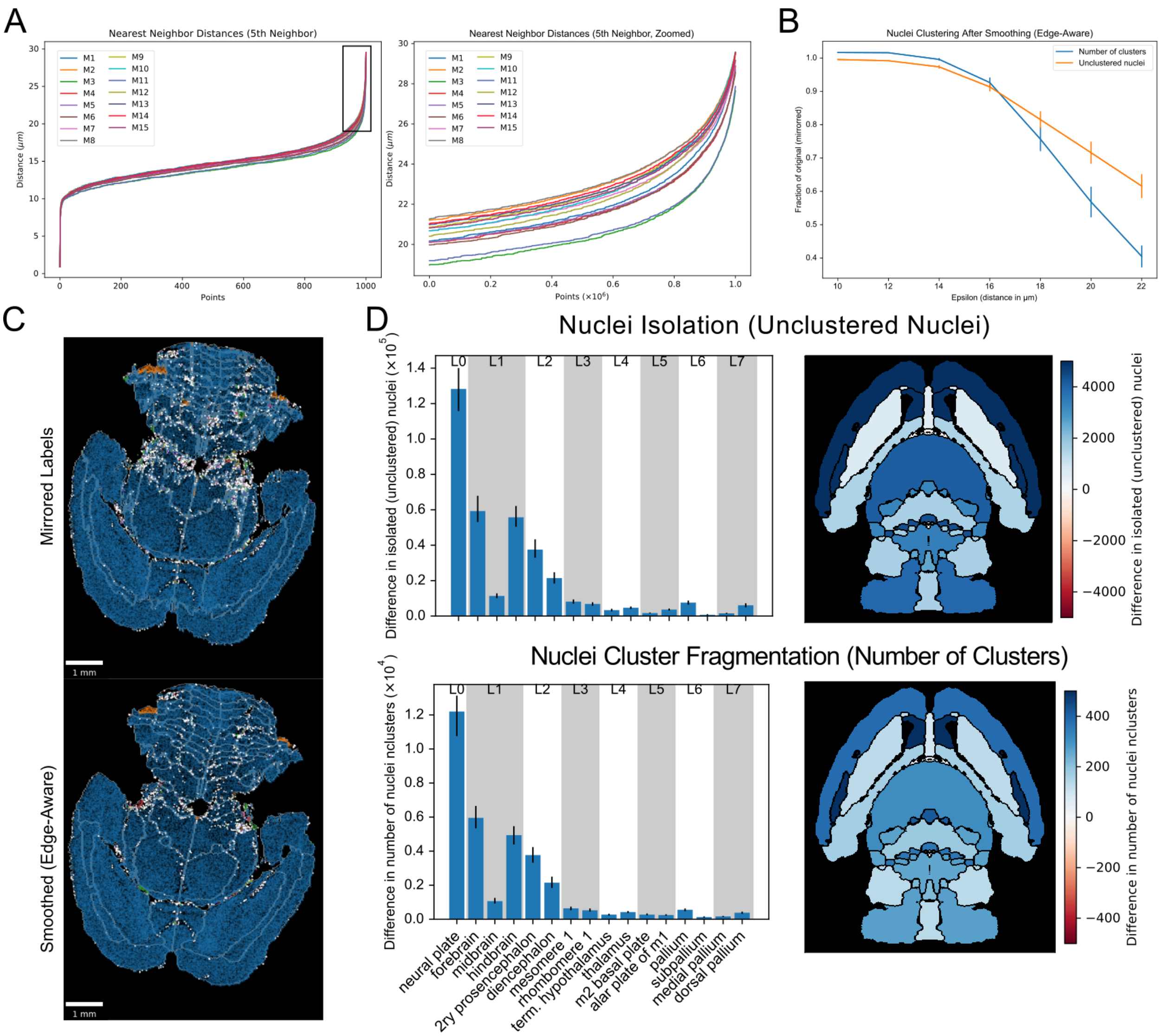


Figure S14: **Nuclei clustering in the original vs. refined atlases** (A) To determine the appropriate cluster neighborhood distance parameter, the fifth nearest neighbor distances for all nuclei in each wild-type brain are plotted in ascending order (right), following the convention in DBSCAN of taking the $2 \cdot ndim - 1$th neighbor [106]. (left) The zoomed view allows identification of the distance at the "elbow" or "knee," or maximum curvature, typically at 20 $\mu m$ or higher. (C) Nuclei clustering within each label in the smoothed (edge-aware) atlas as a fraction of that of the original (mirrored) atlas across a range of distance values. The number of isolated, unclustered nuclei and total clusters, indicative of nuclei isolation and cluster fragmentation, respectively, decreased in the smoothed atlas relative to the original atlas, starting even prior to the minimum distance based on the max curvature seen in part "b." (E) Example using a conservative neighbor distance of 20 for clustering in a brain before (top) and after (bottom) label refinement. Light grey lines show label boundaries. Unclustered nuclei are depicted as white points. All other points are clustered nuclei. Within each region, nuclei are colored by cluster size, where blue represents nuclei in the largest cluster, followed by orange, green, and red for the next successively smaller clusters. (D) Differences by region across all wild-type brains with this distance setting for selected regions across label hierarchies in bar plots (left) and as color gradients on an anatomical map (right).

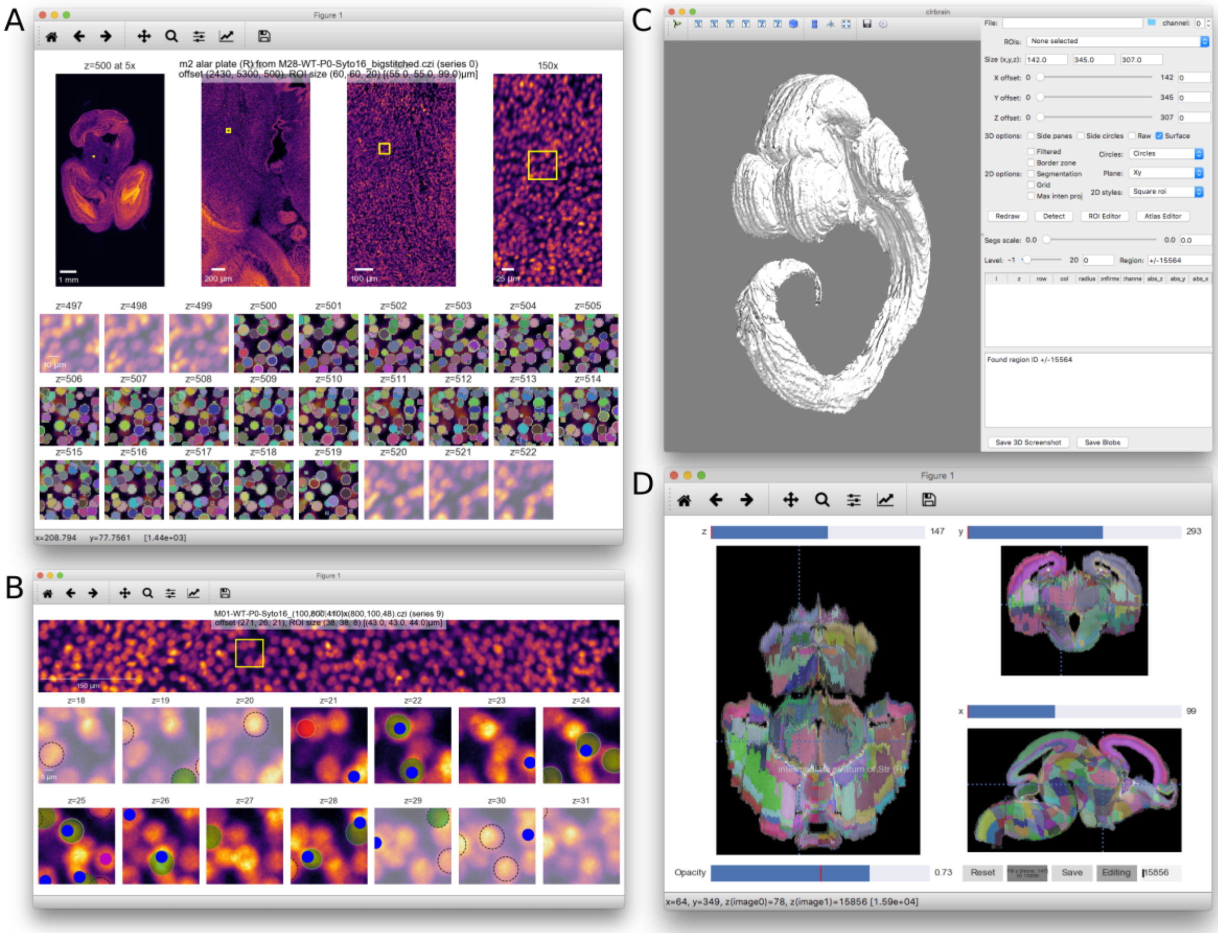


Figure S15: **MagellanMapper software GUI screenshots** (A) ROI serial 2D viewer and annotation editor GUI. The viewer provides overview images at increasing magnifications, zooming into the ROI outlined in yellow. Overview images are scrollable to show how the z-planes change through the ROI in place. Sequential z-planes of the ROI alone depict the original 2D images side-by-side through the entire 3D ROI, with a few additional opacified planes shown above and below the ROI for context. Overlaid labels show an example of automated nuclei segmentation. (B) Another example of the ROI viewer, here in nuclei annotation and verification mode. Automated nuclei centroid "blob detections" are depicted as interactive circles corresponding to blob positions and radii. For building truth sets, circles provide drag-n-drop and copy-paste controls to reposition, add, or subtract these detections for accuracy. Green and red flags allow scoring for detection correctness. To compare with truth sets, blue and purple dots depict previously annotated nuclei positions pulled in from a database. Automated verification of current blob detections against the truth set shows blue dots as correctly detected truth blobs, purple as missed truth blobs, green circles as correct detections based on a matched truth blob, and red as incorrect detections (no remaining truth blob matches). (C) ROI selector and 3D visualization GUI. The right panel displays ROI offset and size controls, 2D and 3D display options, label controls including ontology depth, and an editable blob table. The left panel shows an example whole atlas 3D visualization using the ADMBA E11.5 atlas. (D) Atlas editor GUI. The GUI provides simultaneous orthogonal views in all three dimensions, with planes corresponding to the crosshairs. With "Editing" selected, labels can be painted into adjacent labels as shown in the intermediate stratum of Str label in the left hemisphere of the z-plane. Editing a second, non-contiguous z-plane in the same label will enable the "Fill" button to perform edge interpolation for this label through all intervening planes.

Movie S1: **Original ADMBA E11.5 Atlas**

Movie S2: **3D reconstructed ADMBA E11.5 Atlas**

Movie S3: **Original ADMBA E13.5 Atlas**

Movie S4: **3D reconstructed ADMBA E13.5 Atlas**

Movie S5: **Original ADMBA E15.5 Atlas**

Movie S6: **3D reconstructed ADMBA E15.5 Atlas**

Movie S7: **Original ADMBA E18.5 Atlas**

Movie S8: **3D reconstructed ADMBA E18.5 Atlas**

Movie S9: **Original ADMBA P4 Atlas**

Movie S10: **3D reconstructed ADMBA P4 Atlas**

Movie S11: **Original ADMBA P14 Atlas**

Movie S12: **3D reconstructed ADMBA P14 Atlas**

Movie S13: **Original ADMBA P28 Atlas**

Movie S14: **3D reconstructed ADMBA P28 Atlas**

Movie S15: **Original ADMBA P56 Atlas**

Movie S16: **3D reconstructed ADMBA P56 Atlas**

Data S1: **Spreadsheet mapping colors to label names and IDs in the Allen ontology for each atlas in the ADMBA**

Data S2: **Spreadsheet of metric differences by label for WT brains before and after edge-aware atlas refinement**
